# *Helicobacter pylori* vacuolating cytotoxin A exploits human endosomes for intracellular activation

**DOI:** 10.1101/2022.08.22.504206

**Authors:** Samuel L Palframan, Md. Toslim Mahmud, Kher Shing Tan, Rhys Grinter, Vicky Xin, Rhys A Dunstan, Diana Micati, Genevieve Kerr, Paul J McMurrick, Andrew Smith, Helen Abud, Thanh Ngoc Nguyen, Michael Lazarou, Oded Kleifeld, Trevor Lithgow, Timothy L Cover, Kipros Gabriel, Rebecca J Gorrell, Terry Kwok

## Abstract

*Helicobacter pylori* infection is the main cause of gastric cancer. Vacuolating cytotoxin A (VacA) is a *H. pylori* pore-forming toxin and a key determinant of gastric cancer risk. VacA is secreted as an 88-kDa polypeptide (p88) that upon interaction with host cells induces cytotoxic effects, including cell vacuolation and mitochondrial dysfunction. These effects are currently believed to be due to VacA p88 accumulating inside host cells and forming oligomeric anion-specific channels in membranes of intracellular compartments. However, the molecular nature of intracellular VacA channels in host cells remains undefined. Here we show that VacA p88 does not accumulate inside human epithelial cells, but instead is rapidly processed in endosomes into smaller p31/p28 and p37 products in a manner that precedes VacA-induced vacuolation. VacA processing requires endosomal acidification and concerted cleavage by multiple endo-lysosomal proteases including cathepsins. *In situ* structural mapping reveals that upon processing, the toxin’s central hydrophilic linker and globular C-terminus are excised, whereas oligomerization determinants are retained. Congruently, the processed products are constituents of a high-molecular-weight complex inside the host cell ─ which we propose is the intracellular, mature and active VacA pore. These findings suggest that VacA exploits human endosomes for proteolytic processing and intracellular activation.

**Significance Statement:** *Helicobacter pylori* is a cancer-causing bacterium that infects the stomach of billions of people worldwide. Vacuolating cytotoxin A (VacA) is an important *H. pylori* virulence factor and its activity directly correlates with gastric carcinogenesis. Yet despite decades of intense research, the mechanisms underlying VacA activity in human cells remain incompletely understood. Here, we present evidence suggesting that VacA is activated inside human cells by multi-step proteolytic processing involving endo-lysosomal proteases including cathepsins. We also track and identify the functional processed VacA isoforms in host cells. These results revolutionize our understanding of the mechanism of VacA activation in human cells, whilst expanding our knowledge of the diversity of microbial virulence factors that exploit human endo-lysosomes for pathogenesis.

## Introduction

*Helicobacter pylori* infection of the human stomach is the strongest risk factor for gastric cancer, the fourth leading cause of cancer-related death in the world (1, 2). In the face of decreasing efficacy of *H. pylori* eradication treatment due to burgeoning antibiotic resistance, WHO listed *H. pylori* among 16 antibiotic-resistant bacteria that pose the greatest threat to human health (3).

Vacuolating cytotoxin A (VacA) is a key *H. pylori* virulence factor and a drug and vaccine target involved in *H. pylori* colonization, persistence and pathogenesis (4-7). The 88-kDa mature secreted VacA (designated p88) functions as a pore-forming toxin, but its primary and tertiary structure bear no similarity to other bacterial pore-forming toxins (8, 9). VacA is internalized by host epithelial cells via endocytosis (10) and acts intracellularly to induce cytoplasmic vacuolation (11, 12), mitochondrial dysfunction and endosomal trafficking perturbation (13-15). A large body of epidemiological data indicate that VacA channel activity correlates with increased disease severity and a heightened gastric cancer risk (16).

Extensive biophysical studies show that VacA p88 exhibits weak anion-specific channel conductance in both artificial planar lipid bilayers and plasma membranes of intact epithelial cells (14, 17, 18). Channel activity is strictly dependent on the ability of VacA to self-associate into higher-order oligomers (19-21). Secreted VacA p88 consists of two domains, an N-terminal p33 domain and a C-terminal p55 domain, which are both essential to pore-forming activity and receptor-binding (13). The two domains are connected by a central hydrophilic linker that is dispensable for anion channel activity and prone to proteolytic-nicking *in vitro* (22-24). Recent high-resolution cryo-EM structures of water-soluble VacA p88 oligomers further suggests that this hydrophilic linker, likely located at the center of the flower-shaped VacA p88 hexamer (Fig. 1), is highly flexible in structure (8). It was previously speculated that VacA resembled classical A-B toxins and might undergo proteolytic cleavage within the hydrophilic linker inside host cells (25). Nevertheless, intracellularly processed functional VacA domains have not been reported to date. Meanwhile, a widely accepted model of VacA mechanism of action proposes that the p88 hexamer forms the active channel responsible for most of the toxin’s cellular effects (18). The physiological significance of this model has not yet been rigorously examined.

**Figure 1.**
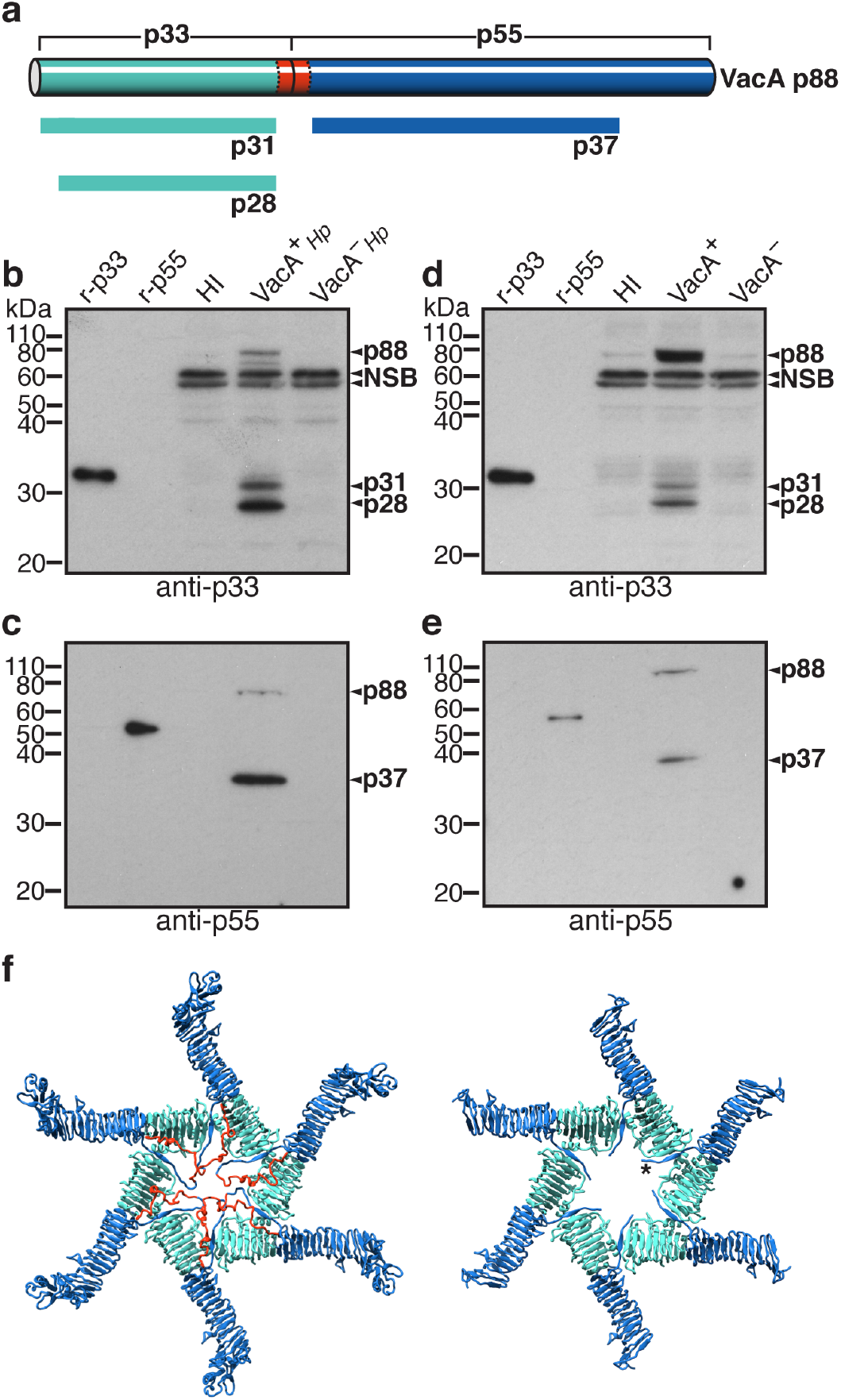
Detection of novel N- and C-terminal VacA cleavage products in *H. pylori*-infected AGS cells and VacA-intoxicated HeLa cells. **a**, Summary of novel VacA cleavage products detected in human cells; top cylindrical bar represents VacA p88, hydrophilic linker in red; novel VacA forms represented by rectangular bars. Immunoblot analysis of lysates from AGS cells co-cultured with VacA^+^*Hp*, VacA^−^*Hp* or heart infusion (HI) broth for 24 h using **b**, anti-p33 or **c**, anti-p55. Immunoblot analysis of lysates from HeLa cells treated with CS from VacA^+^*Hp* (VacA^+^), VacA^‒^ *Hp* (VacA^‒^) or HI broth for 24 h using **d**, anti-p33 or **e**, anti-p55; NSB = non-specific band. **f**, Cryo-EM structure of VacA p88 hexamer (8) (left; PDB ID: 6NYF) and novel VacA fragments p31/p28 and p37 modelled on same structure (right) using UCSF Chimera (61); *C-terminal domain segment that cradles adjacent protomer; p33 domain and derived fragments (cyan), p55 domain and derived fragment (blue), hydrophilic linker region (red) as predicted by SWISS-MODEL software (58).

Despite VacA acting intracellularly as a pore-forming toxin (11), knowledge of the physiological form inside the host cell is lacking. In this study, we directly address this key knowledge gap using an *in situ* structural mapping approach. The data provide crucial insights into the physiological form of the VacA channel in human epithelial cells. Our findings suggest that VacA p88 undergoes rapid proteolytic modification and activation in the endosome of host cells prior to inducing vacuolation.

## Results

### VacA is processed by human epithelial cells

To gain insight into the physiological form of VacA inside human gastric epithelium, we developed domain-specific antisera capable of recognizing N-terminal p33 or C-terminal p55 domain regions of VacA (*SI Appendix*, Fig. S1a,b). Using these antisera, we examined the steady-state levels of VacA in a human gastric adenocarcinoma (AGS) cell line infected with *H. pylori* 60190wt (VacA^+^*Hp*). In addition to cell-associated VacA p88, we detected novel forms of VacA as summarized (Fig. 1a): the antiserum raised to the N-terminal p33 domain detected distinct proteins that migrated at 31 kDa (p31) and 28 kDa (p28) (Fig. 1b), whereas the antiserum raised to the C-terminal p55 domain detected a distinct protein that migrated at 37 kDa (p37) (Fig. 1c). These proteins were derived from VacA p88 since they were absent from cells inoculated with a *H. pylori* 60190 VacA-null mutant (VacA^‒^*Hp*) or heart infusion broth (HI) culture media alone (Fig. 1b,c). The observed processing of VacA was not limited to the AGS cell line but is likely a fundamental process since HeLa cells similarly treated with VacA-containing culture supernatant (CS) showed the same processed forms of VacA (Fig. 1d,e).

Multiple lines of evidence confirm that these VacA fragments are generated by precise intracellular processing and are not degradation products. First, even after 24 h incubation with AGS cells, the only antisera-reactive band detected in spent culture medium recovered from VacA-treated cells was p88 (*SI Appendix*, Fig. S2a). This indicated that p28, p31 and p37 do not arise from extracellular degradation of p88. Second, endocytosis inhibitors cytochalasin D and EGA (26) that blocked uptake of VacA also prevented the generation of p31, p28 and p37 (*SI Appendix*, Fig. S2b,c), suggesting that these fragments arise only after VacA entry into the host cell. Third, VacA processing in AGS cells treated with proteasome inhibitor MG132 was indistinguishable from that in untreated cells; this implies that proteasomal degradation does not contribute to the formation of p28, p31 and p37 (*SI Appendix*, Fig. S3a). Fourth, Atg8 family Hexa KO HeLa cells, which lack the six known Atg8s and are therefore deficient in autophagy (27), processed VacA identically to wild-type HeLa cells (*SI Appendix*, Fig. S3b), confirming that autophagy does not contribute to VacA processing.

Finally, the physiological relevance of VacA processing was confirmed by the fact that similar processed fragments were observed in VacA-intoxicated human two-dimensional (2D) primary gastric organoid-derived monolayers (*SI Appendix*, Fig. S4). Taken together, these observations strongly suggest that VacA is processed inside host cells into hitherto unknown fragments by intracellular proteases.

### VacA processing correlates with function

Given that many intracellularly-acting bacterial toxins are activated inside the host cell (28), we next investigated whether the VacA p28/p31 and p37 fragments are associated with vacuolating activities. First, we assessed the kinetics of VacA processing and its relationship to vacuolation using a cumulative intoxication assay. By treating AGS cells for up to 6 h with VacA^+^, we found that both p31 and p37 were detectable as early as 1 h post-intoxication (*SI Appendix*, Fig. S5a). Their emergence preceded the onset of vacuolation, as assayed by neutral red uptake (NRU) (*SI Appendix*, Fig. S5c), and both fragments continued to accumulate over time. The accumulation of p28 started slightly later, at ∼4 h post-intoxication (*SI Appendix*, Fig. S5a). More importantly, experiments employing trypsin-shaving of VacA-intoxicated cells showed that VacA p88 accumulated only on the cell surface and not inside host cells (*SI Appendix*, Fig. S5b). The most plausible explanation for these observations is that endocytosed VacA p88 is processed rapidly into specific fragments and hence does not accumulate inside the host cell. Thus, contrary to longstanding belief, these observations argue that VacA hexamers composed of p88 are unlikely to represent the intracellular physiological VacA channel, but rather a prepore state.

To dissect the sequence of VacA processing and assess whether these processed products correspond with intracellular activities, we analyzed VacA-intoxicated cells in pulse-chase experiments. AGS cells were pulsed with VacA^+^ for 30 mins, after which unbound toxin (p88) was removed. Cell-associated VacA was then examined over time (up to 10 h post-pulse) by immunoblot analysis of intoxicated-cell lysates using p33- or p55-specific antisera (Fig. 2a). Consistent with our previous observations of negligible intracellular p88, increasing p31 and p37 levels directly correlated with declining p88 levels (Fig. 2b,c, respectively). Additionally, p28 became detectable from 2 h post-pulse, with its accumulation coinciding with the loss of p31 over time (Fig. 2b), suggesting a two-tiered processing of VacA in which p31 is further processed to yield p28. By 8 h and 10 h post-pulse, p28 and p37 were the predominant forms of intracellular VacA.

**Figure 2.**
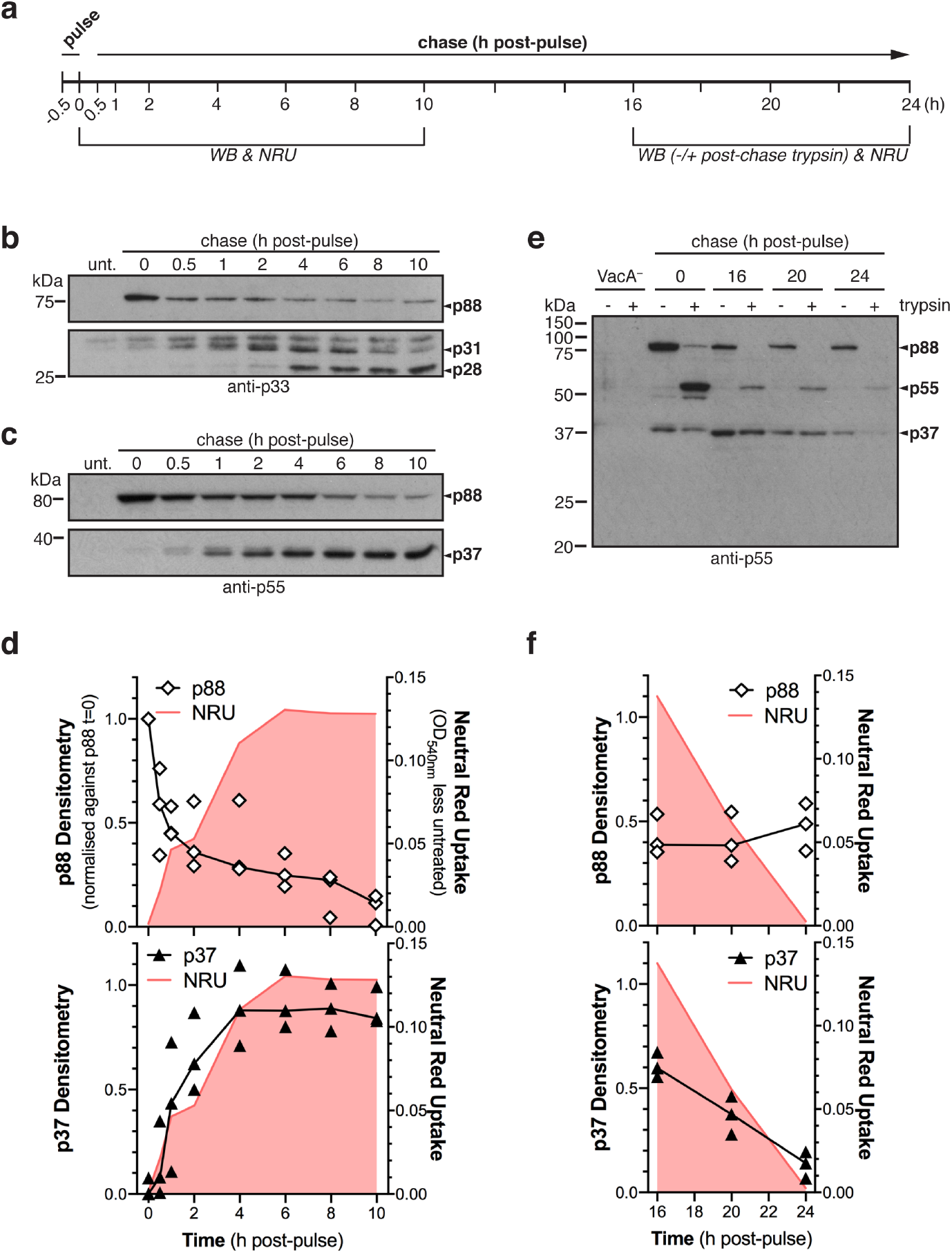
VacA p88 is processed into active fragments over time by host cells. VacA^+^ pulse-chase time-courses in AGS cells **a**, timeline of pulse-chase experiments. **b**, Immunoblot analysis of AGS lysate from 0 - 10 h pulse-chase using anti-p33 or **c**, anti-p55. **d**, Densitometry analysis (N=3 independent experiments) of 0 - 10 h pulse-chase in AGS cells overlaid on parallel neutral red uptake (NRU) data. **e**, Immunoblot analysis (anti-p55) of AGS lysate from 16 - 24 h pulse-chase without (-) or with (+) trypsin-shaving at harvest. **f**, Densitometry analysis (N=3 independent experiments) of 16 - 24 h pulse-chase in AGS cells overlaid on parallel NRU data. NH_4_Cl was added to cell culture media 0.5 h before harvest to potentiate vacuolation (panels b–f); upper and lower sections of panel b were from separate blots prepared using the same samples; anti-p55 immunoblots used for densitometry; lines in densitometry and NRU plots denote median of three independent experiments; see *SI Appendix*, Figure S6a for individual data points and statistical analysis of NRU data.

Strikingly, upon NRU analysis of a parallel set of identical VacA-pulsed AGS monolayers (*SI Appendix*, Fig. S6a), we observed that VacA-mediated NRU was inversely proportional to p88 levels (Spearman r = −0.91, p = 0.005) but positively correlated with increasing levels of processed VacA (Spearman r = 0.93, p = 0.002) (Fig. 2d; *SI Appendix*, Fig. S6b). This suggests that processed VacA, rather than full-length p88, is responsible for vacuolation. To further examine this hypothesis, we tested whether depletion of processed VacA would lead to loss of vacuolating activity. For this, we pulsed cells with VacA^+^ and then sampled cell lysates for immunoblot analysis (anti-p55) and NRU assay at the later time points of 16, 20 and 24 h post-pulse, thereby allowing for natural depletion of intracellular VacA (Fig. 2a). An identical replicate set of VacA^+^-pulsed AGS cells were trypsin-shaved immediately before harvest to identify residual cell surface-bound p88 by cleaving it into p33 and p55, and analyzed by anti-p55 immunoblot (Fig. 2e). These results showed that the gradual loss of vacuolating activity after 16 h mirrored the decreasing p37 levels, but did not correlate with p88 levels, which was shown to be entirely extracellular by trypsin-shaving (Fig. 2e,f). Collectively, the findings from these pulse-chase experiments support a new model for VacA mechanism of action: VacA p88, upon entry into host cells, is rapidly processed into p31/p28 and p37 fragments that trigger robust vacuolation until their clearance from the cell.

### Fragments associate with early endosomes and form a protein complex

Since VacA exerts its vacuolating activity by functioning as an oligomeric anion channel in the endosomal membrane of host cells (10, 14, 19-21, 29, 30), it is thus tempting to postulate that the processed VacA fragments p31/p28 and p37 exist as an oligomeric complex in the host endosome. To address this hypothesis, we first examined the subcellular localization of p31, p28 and p37 using sucrose density gradient centrifugation. Analysis of the AGS cell extracts showed that the processed VacA products p31, p28 and p37 cofractionated substantially with the early endosomal marker Rab5 (Fig. 3a), suggesting their localization in early endosomes (fractions 4–6) (10). In addition, p31 and p37 were also evident in the most prominent mitochondrial fractions (fractions 7,8). Mitochondrial outer membrane protein Sam50 cofractionated not only with the VacA processed fragments but also Rab5, whereas cells treated with VacA^−^ (Fig. 3a) displayed less Rab5-Sam50 co-fractionation, observations that are consistent with a previous study demonstrating that VacA promotes endosome-mitochondria juxtaposition (31). Together, these results support a model where p28, p31 and p37 accumulate within endosomes and mitochondria-associated endosomes.

**Figure 3.**
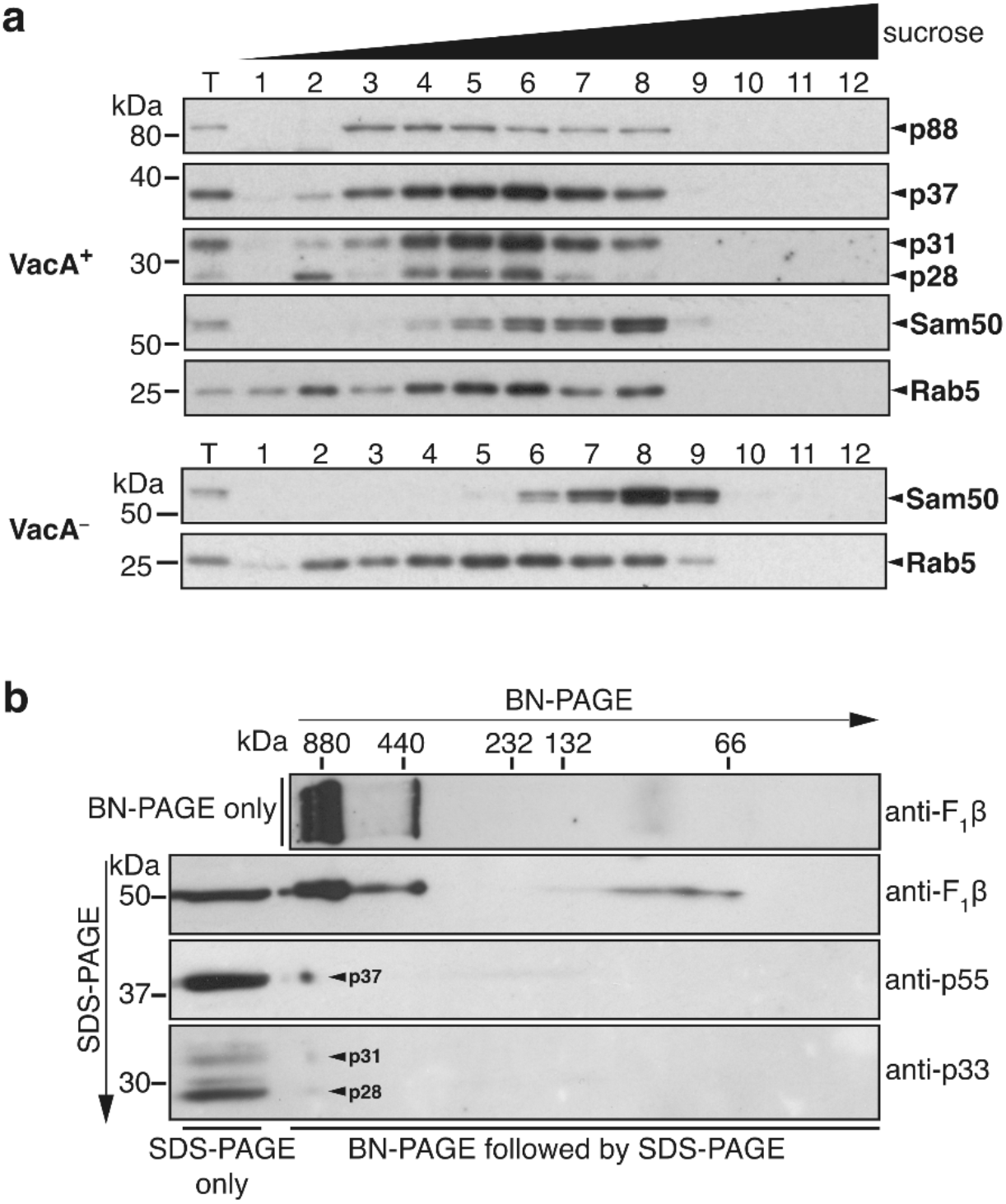
VacA processed products associate with early endosomes and are constituents of a higher-order complex. **a**, Immunoblot of sucrose gradient-fractionated VacA^+^- or VacA^−^-treated (6 h) AGS cells; VacA (anti-p55 for p88/p37; anti-p33 for p31/p28), mitochondria (Sam50) or early endosome (Rab5) antibodies; T = total cell lysate. **b**, 2D-PAGE and immunoblot analysis of endosomal/mitochondrial-enriched fraction of lysed AGS cells following VacA^+^ treatment (24 h) using the indicated antibodies; F_1_β is the beta subunit of F_1_-ATPase complex in mitochondria, present in ∼1000 kDa F_1_F_0_ dimer, ∼700 kDa F_1_F_0_ monomer and ∼400 kDa F_1_ portion of the complex when solubilized in digitonin (62) and serves as a BN-PAGE control.

Oligomerization is crucial for VacA to function as anion channels both *in vitro* and inside host cells (21, 32). We thus next interrogated whether p31/p28 and p37 co-exist as oligomers in subcellular membranes using a 2D-PAGE approach. Employing non-denaturing BN-PAGE followed by SDS-PAGE, we analyzed an endosome- and mitochondria-enriched fraction of VacA-treated AGS cells. The data showed that subcellular organelle-derived VacA co-migrated as a protein complex containing p31, p28 and p37 (Fig. 3b; *SI Appendix*, Fig. S7). Taking into account the exaggerating impact of detergent/dye binding on membrane protein size estimation in BN-PAGE (33), the results suggest that these complexes are hexameric, or higher-order oligomers that represent VacA anion channels in subcellular membranes.

### Fragments retain crucial VacA regions

To further elucidate the structural nature of the intracellular processed VacA, we employed mass spectrometry (MS) analysis to determine the amino acid sequences of the various VacA processed fragments. Tryptic/chymotryptic peptides yielded from p31 and p28 span amino acid residues G31 to K274, indicating that both fragments contain β-strands 1–15 in the N-terminal p33 domain of VacA (8) (Fig. 1a; *SI Appendix*, Fig. S8a; Table S1). By comparing the intracellular processing of wild-type VacA with VacA^Δ6-27^, we discovered that p31 was replaced by a truncated form in cells infected with the VacA^Δ6-27^ strain, whereas wild-type VacA and VacA^Δ6-27^ both generated identical p28 forms (*SI Appendix*, Fig. S8b). This reveals that the difference between p31 and p28 is the presence and absence, respectively, of the VacA N-terminus (residues A1-W30), a key hydrophobic region involved in channel formation (34). Given these results and the estimated molecular weight of the fragments, both p31 and p28 likely terminate at the beginning of the inter-domain hydrophilic linker, specifically at or around P294 (Fig. 1a).

Tryptic/chymotryptic peptides from VacA p37 spanned residues T337 to K689 of VacA p88, corresponding to β-strands 1–29 in the C-terminal p55 domain (8) (*SI Appendix*, Fig. S8a; Table S1), and p37 peptides containing a ragged N-terminus indicated that V339 is the N-terminal boundary of p37 (*SI Appendix*, Table S1). Thus, p37 contains the V339-A350 segment that is essential for oligomer formation (30, 35) via cradling of the p33 domain of the adjacent protomer, as shown in the cryo-EM structure of p88 oligomers (8, 9) (Fig. 1f). Collectively, these results reveal that the VacA processed fragments p31, p28 and p37 contain crucial functional and structural determinants, and that both the central hydrophilic linker and the C-terminal ‘foot’ region are excised during VacA processing (Fig. 1f).

### Processing involves endosomal acidification and cathepsins

Multiple disparate VacA cleavage sites together with the rapid p88-turnover in host cells suggests that intracellular VacA processing occurs via a series of robust cleavage events. Many intracellularly-acting bacterial toxins and viral fusion proteins exploit endosomal acidification and endo-lysosomal proteases for proteolytic processing (36-38). To first assess whether VacA processing requires endosomal acidification, AGS cells were treated prior to and during VacA^+^ intoxication with the endosomal acidification inhibitors bafilomycin A1 (vacuolar-H^+^-ATPase inhibitor) or chloroquine (acidotropic weak base). These inhibitors blocked p31/p28 and p37 production (Fig. 4a,b), resulting in p88 accumulation inside the intoxicated host cells as evidenced by the protection of a proportion of p88 from exogenous trypsin digestion (Fig. 4a,b). Furthermore, treatment of cells with the acidotropic weak base ammonium chloride, commonly used to potentiate VacA-induced vacuolation, slowed p88 processing in VacA-intoxicated cells (*SI Appendix*, Fig. S9). This is consistent with observations of ammonium chloride-enhanced p88 intracellular persistence previously reported by Foegeding et al. (39). Taken together, our findings that endosomal acidification inhibitors block intracellular p88 processing into p31/p28 and p37 suggest that endosomal acidification is essential for intracellular processing of p88. Interestingly, VacA processing in HeLa Vps39 knockout cells (Vps39 KO), which lack a gene product essential for endosomal maturation (27), yielded an additional band at ∼47 kDa and a doublet at ∼38-39 kDa, but no p37 (*SI Appendix*, Fig. S3b); this confirms the central role of the endosomal pathway in host-cell mediated VacA processing.

**Figure 4.**
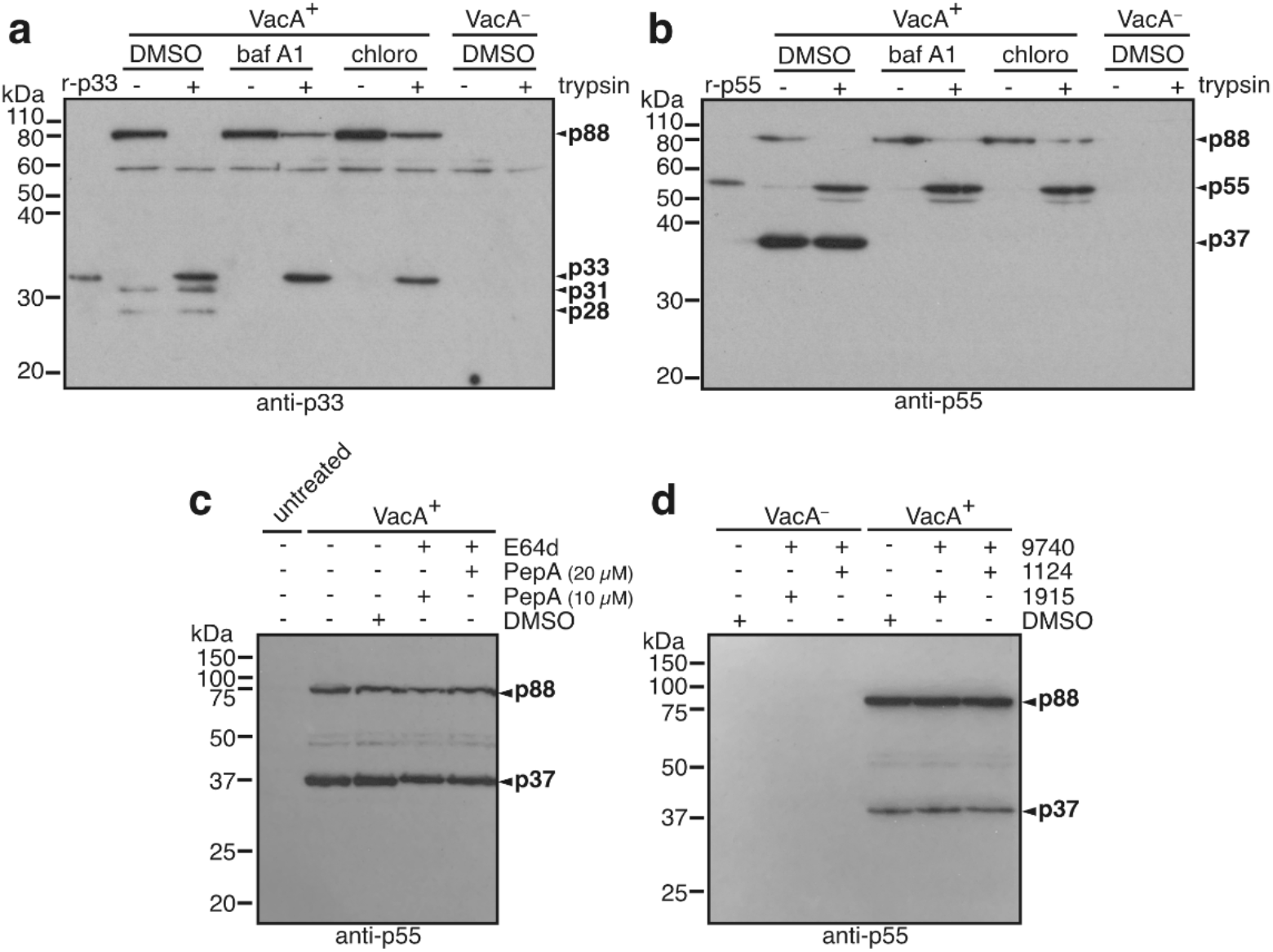
VacA processing requires endosomal acidification and involves cathepsin proteases. **a**, Immunoblot analysis using **a**, anti-p33 or **b**, anti-p55 of VacA^+^-treated AGS cells without (DMSO) or with endosomal acidification inhibitor (bafilomycin A1 or chloroquine) before harvest without (-) or with (+) trypsin. Immunoblot analysis using anti-p55 of VacA^+^-treated AGS cells without (DMSO) or with **c**, Pepstatin methy ester (PepA) which is an aspartic protease inhibitor and the cysteine protease inhibitor E64d as indicated; **d**, cathepsin S inhibitors (1915, 1124) and cathepsin C inhibitor (9740) as indicated.

To further understand the molecular mechanism behind VacA processing, we used a panel of specific cathepsin inhibitors. The cathepsin family of proteases represent the main class of acid hydrolases in endosomes and lysosomes. Pre-treatment of AGS cells with Pepstatin A methyl ester (a membrane-permeable cathepsin D and E inhibitor) and E64d (membrane-permeable and highly specific cathepsin B, H, L and K inhibitor) yielded a p55-reactive processed fragment slightly larger than the p37 fragment (Fig. 4c), which was not observed when cells were treated with Pepstatin A methyl ester alone (data not shown). These data suggest that cathepsins B, H, L and/or K is/are involved in VacA processing. In contrast, cathepsins C and S do not seem to be required as pretreatment with their corresponding specific inhibitors did not alter the cleavage pattern (Fig. 4d). The fact that none of these protease inhibitors, either alone or in combination, resulted in complete blockade of VacA processing or vacuolation (data not shown) suggests that VacA processing is a robust and complex process involving multiple endo-lysosomal proteases. This is perhaps unsurprising given the proteolytic susceptibility of the flexible hydrophilic linker in VacA and the broad-spectrum substrate specificities of cathepsins.

## Discussion

VacA is a key virulence factor of *H. pylori* and an important disease marker. The current understanding of its structure and function is largely based on extensive research on water-soluble VacA hexamers *in vitro*. It is speculated that the membrane-associated active VacA channel might be similar in structure to water-soluble VacA hexamers (8, 40), of which two high-resolution cryo-EM structures were recently published (8, 9, 40). Despite these reasonable assumptions, the actual nature of the physiological intracellular anion channel has remained unknown since the discovery of VacA over thirty years ago (41-43). Here, our data provide crucial insights into the intracellular active form of VacA and reveal a new paradigm of a bacterial pore-forming toxin that exploits the human endosome for spatiotemporal activation. Our findings suggest that the hydrophilic linker and globular C-terminus are excised from full-length VacA to generate functional N-terminal p31/p28 and C-terminal p37 fragments in endosomes following endocytosis.

A key discovery in this study is that the only VacA species accumulating in the endosome are the processed VacA fragments and not full length VacA. It is well established that vacuolation is caused by VacA functioning as anion channels in the endosomal membrane (14, 44). Thus, this novel finding has crucial implications because together with the strict temporal correlation of VacA processing with vacuole development and resolution, it argues unequivocally that the processed form, and not full length unprocessed VacA, is the active VacA form that causes vacuolation in the endosome.

The hydrophilic linker that separates the N- and C-terminal VacA domains is vulnerable to *in vitro* proteolytic nicking (22, 23, 45) and is dispensable for anion channel activity (24), which led to speculation that VacA might represent a prototypical A-B type pore-forming toxin (25, 46). Recent cryo-EM structures of water-soluble hexamers predicted that the hydrophilic linker occupies the central cavity of oligomers (8), in close proximity to an N-terminal glycine zipper-containing hydrophobic α-helix, which is essential for insertion of the functional channel into membranes (19, 34). Excision of the linker, as reported in this study, may represent the first step of VacA channel activation, whereby the N-terminal glycine zipper is dissociated from the linker facilitating membrane insertion following endocytosis (Fig. 5). Meanwhile, cleavage of the ‘foot’ in the globular C-terminus of VacA, speculated to be involved in host cell binding (47), may help liberate VacA from bound host cell receptors within the endosome. Such molecular workings may be a crucial mechanism to ensure full channel activation only in intended environments, such as endosomes, and may serve to avoid non-specific membrane damage.

**Figure 5.**
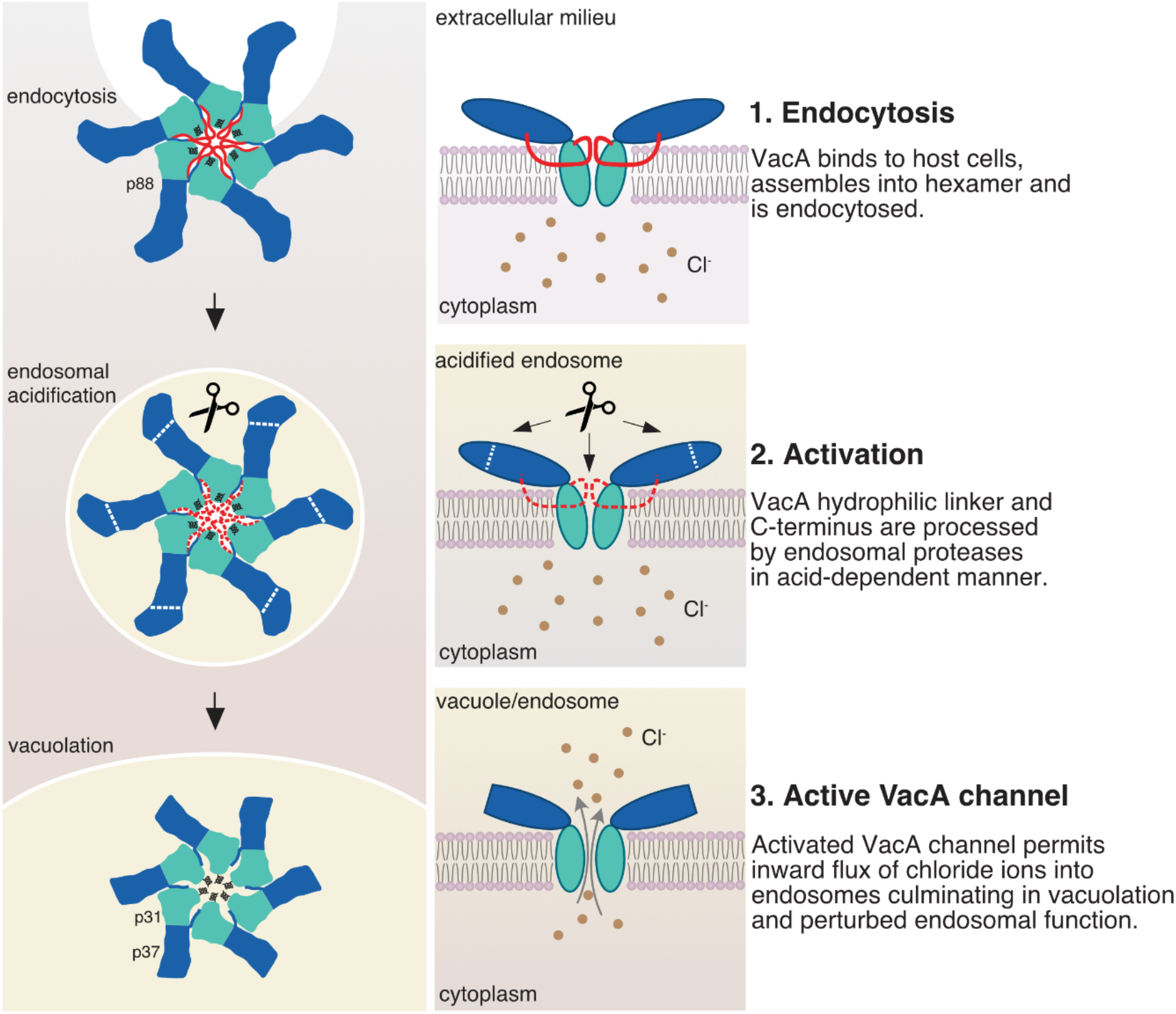
Proposed model for VacA activation inside human endosomes: a new VacA intoxication paradigm. **1)** The VacA hexamer is endocytosed by host cells; **2)** Endosomal acidification induces conformational change in the VacA oligomer that exposes previously protected residues to early endosomal proteases. The hydrophilic linker (red) and C-terminal ‘foot’ are then proteolytically excised from p88 to generate p31 and p37 VacA fragments; **3)** Excision of the linker may enhance mobility of the hydrophobic N-terminus due to their predicted close proximity in the centre of the assembled oligomer (8), thereby initiating membrane insertion and formation of the active channel. Notably, the beta-sheet extension of the p55 domain that cradles adjacent protomers is retained in fragment p37 and may be critical to stabilize processed VacA oligomers. The processed and active VacA oligomer dissipates the established ion concentration gradient between endosomes and cytoplasm culminating in vacuolation. Additional proteases present in late endosomes/endo-lysosomes cleave the hydrophobic N-terminus of p31 to generate p28, which likely also possesses channel activity. The N-terminal and C-terminal structural domains of VacA are shown in cyan and blue, respectively.

Our findings clarify various longstanding unsettled issues related to the intracellular form of VacA. Previous studies based on ectopic expression of VacA constructs in HeLa cells suggest that the first 422 amino acid residues of VacA suffice to cause vacuolation (12), but the implications of these observations remained unclear. In this study, our results suggest that VacA is processed within host cells to generate functional N-terminal p31/p28 and C-terminal p37 fragments. Notably, these fragments together comprise the 422 amino acid residues in the aforementioned minimal vacuolation-competent VacA construct (12) minus the central hydrophilic linker (24). Thus, our findings provide a plausible mechanistic explanation for these previously puzzling observations. Our findings also shed light on a twenty-year-old enigma in the field surrounding the ability of VacA to cause vacuolation despite deletion or substitution of the entire hydrophilic linker (24). Our discovery that the hydrophilic linker is in fact excised during VacA proteolytic activation provides a logical explanation for these once intriguing observations. Finally, while some studies provided evidence that regions from both N- and C-terminal VacA domains were required for vacuolation (48, 49), one study showed that the N-terminal domain alone, when chemically crosslinked *in vitro*, was sufficient for anion channel activity (50). The domain-specific VacA antibodies developed in this study have enabled us to clarify this discrepancy. Our data show that regions from both the N- and C-terminal domains comprise the intracellular and active VacA channel, with the premise that the N-terminal p31/p28 processed fragments constitute the channel portion of the mature VacA pore, whereas the C-terminal p37 fragment plays a structural role to stabilize the oligomeric pore complex. The latter is in accordance with recent cryo-EM structural data and data from other studies showing that residues V339-A350, which are retained in fragment p37, are important for toxin oligomerization and stability (8, 30).

There are two caveats in our findings that have implications for the complex intracellular mechanism of action of VacA. One concerns the observed sequential processing of p31 to p28 with an apparent loss of the N-terminal hydrophobic α-helix. Based on previous data (19), one would expect loss of this helix to result in a non-functional VacA toxin. However, our pulse chase analyses suggest that p28, like p31, is functional. This apparent paradox could be explained if the hydrophobic N-terminal α-helix of VacA is required for membrane insertion and/or the initial formation of a functional channel, but is dispensable once the mature channel has formed in the endosome. Alternatively, in the mature VacA channel, the hydrophobic N-terminal α-helix of VacA may still be present but is no longer covalently-associated with the rest of the protein; this is conceivable if the short linker immediately upstream of hydrophobic N-terminal α-helix would undergo a conformational change in the processed active VacA form that renders the linker more susceptible to proteolytic cleavage. Finally, although we have identified cathepsins B, H, K or L to be involved in host cell processing of VacA, we were unable to completely block processing despite an exhaustive screen of protease inhibitors. This suggests that VacA processing involves multiple proteases and/or that the VacA cleavage regions confer protease-promiscuity and redundancy. These features support the notion that VacA has evolved to undergo robust endosomal processing – which would be expected if the latter is an integral step in the intracellular activation of VacA in endosomes.

VacA has been shown to exhibit weak anion channel activity in the plasma membrane of host cells (17). Our findings are therefore in line with the model that VacA is capable of a ‘dual mode’ operation – the extracellular full length p88 channel is capable of exerting weak channel activity in the plasma membrane, whereas the processed form reported in this study is responsible for the more cytotoxic intracellular channel activities, including vacuolation, that need to be under tight spatiotemporal control.

In summary, this study has redefined the physiological active form of VacA. It has provided data in support of a new model in which VacA exploits human endosomal machinery for intracellular activation. Thus, VacA, despite its unique features and functional properties, is reminiscent of some A-B toxins such as diphtheria toxin and anthrax toxin, whose cellular functions similarly rely on endosomal acidification and proteolytic activation (51). Given the crucial role of VacA in *H. pylori* pathogenesis, future identification of the protease(s) responsible for VacA activation could open doors to therapeutic innovations to combat antibiotic-resistant *H. pylori* infection and the associated gastric cancer.

## Materials and Methods

### Cell and bacterial culture

AGS cells, a human gastric adenocarcinoma cell line, and HeLa cells, including Hexa KO and Vps39 KO mutant HeLa cell lines (*SI Appendix*, Table S4), were grown in Dulbecco’s modified eagle medium (DMEM; Gibco) supplemented with 10% (v/v) fetal bovine serum (FBS; Serana) at 37 °C in a humidified atmosphere containing 5% CO_2_. *H. pylori* strains (*SI Appendix*, Table S4) were routinely maintained and broth cultures prepared as described previously (52).

### Mutant *H. pylori* strains

Donor DNAs for creating *H. pylori vacA* mutant strains were synthesized from *H. pylori* 60190 nucleotide sequence (accession number U05676) and cloned into pUC57 (Genscript) in two overlapping parts: part A contained nucleotides 210 to 1420, with nucleotides 597 to 600 substituted to GGAT to create a BamHI site; part B contained nucleotides 1321 – 2884 and was synthesized as either untagged, or carrying extra nucleotides encoding a monoclonal antibody-recognizable Strep tag II epitope tag (SAWSHPQFEK) inserted after amino acid 234 (p31-tagged) or 525 (p37-tagged). Part A was further modified by ligating a *cat* gene in the created BamHI site. Parts A and B were amplified using primer pairs VacA_NtermF/R or VacA_CtermF/R, respectively, and then part A was joined to one of each of the part B forms using splice-by-overlap PCR using primer pair VacA_NtermF/VacA_CtermR with Phusion DNA polymerase (primers in *SI Appendix*, Table S5). *H. pylori* 60190 VacA-null mutant (VacA^−^*Hp*) was naturally transformed using the spliced PCR products to produce tagged strains 60190 Str_234_ and 60190 Str_525_. Transformants were selected on GC plates supplemented with 8 µg/ml chloramphenicol and screened for vacuolating activity, sensitivity to kanamycin, and insertion of the tag by colony PCR, where relevant.

### *H. pylori* culture supernatant (CS) preparation

Broth cultures of *H. pylori* strains 60190wt, 60190 VacA-null mutant or P12 were inoculated to an O.D_600_ of 0.05 and grown for 24 hours (h). Bacteria were pelleted by centrifugation at 4000 x*g* for 20 min and the supernatant filter-sterilized using Millipore EXPRESS^™^ Plus 0.22-μm membrane filter vacuum stericups. Sterile supernatants were three-times concentrated using Millipore Amicon Ultra-15 centrifugal 3,000 nominal molecular weight limit (NMWL) (Da) cutoff units by centrifuging at 2500 x*g* for ∼40 minutes (min) at 4 °C. HI or brain heart infusion broth (BHI) negative control was prepared in the same manner. Sterile, concentrated VacA^+^, VacA^−^ and P12 CS were stored in aliquots at −80 °C.

### Production of VacA p33Δ6-27 and p55 recombinant proteins and specific polyclonal antisera

Plasmids carrying DNA encoding *H. pylori* 26695 VacA subunits p33Δ6-27 and p55 were amplified using specific primer pairs p33F/p33R and p55F/p55R (*SI Appendix*, Table S5), respectively, and ligated into BamHI/SalI-linearized expression vector pQE-30 (Qiagen) to produce pQE30-p33Δ6-27 and pQE30-p55. N-terminal His-tagged recombinant p33Δ6-27 (r-p33) and p55 (r-p55) proteins were expressed as inclusion bodies in *E. coli* BL21-DE3 and BL21-DE3 Codon Plus, respectively, by inducing a mid-log culture with 0.5 mM IPTG (3 h at 37 °C). Following cell lysis using lysis buffer (0.1 % (v/v) Triton X-100, 0.2 mg/ml lysozyme, 10 µg/ml DNaseI, 0.1 % (v/v) β-mercaptoethanol in PBS, pH 7.5) and homogenization (Avestin cell crusher, 2 passes), inclusion bodies were pelleted (20,000 x*g*, 30 mins, 4 °C) and solubilized overnight at 4 °C in binding buffer (4 M urea, 200 mM NaCl, 20 mM Tris, pH 7.5). His-tagged proteins were purified using Ni-NTA resin (Promega); bound proteins were washed with binding buffer and eluted into elution buffer (4 M urea, 200 mM NaCl, 400 mM imidazole, 20 mM Tris, pH 7.5). Purified proteins were concentrated by ultrafiltration (Amicon Ultra-4, Millipore) and diluted in PBS.

For antisera production, New Zealand White rabbits were immunized with 250 µg recombinant protein in Freunds complete adjuvant, and boosted 4 times with 150 µg in Freunds incomplete adjuvant at 2-4 week intervals. Specificity of polyclonal antisera was confirmed against r-p33 and r-p55 proteins, and native *H. pylori* VacA protein by immunoblot. All animal work was compliant with the Australian National Health and Medical Research Council guidelines and approved by the Monash University School of Biomedical Sciences Animal Ethics Committee.

### Intoxication of cultured mammalian cells with VacA

AGS and HeLa cells for *H. pylori* CS-treatment experiments were grown to ∼80 % confluence before the addition of three-times concentrated *H. pylori* CS to a final concentration of 20 % (v/v) in cell culture media, and incubated at 37 ^°^C for 24 h unless otherwise stated. AGS cells for supernatant time-course experiments were seeded at 7.5 × 10^4^ cells per well in a 24-well plate or 1 × 10^4^ cells per well in a 96-well plate and incubated with *H. pylori* CS the following day for the indicated durations. For pulse-chase experiments, cells were intoxicated for 30 mins with VacA^+^ or VacA^−^ before one wash with PBS to remove unbound toxin, fresh media was replaced in wells for the remainder of the experiment. For *H. pylori* co-culture experiments, AGS cells were seeded at 1 × 10^5^ cells per-well in a 24-well plate and incubated the following day with broth-cultured *H. pylori* strains (moi = 30) or sterile HI broth for 24 h, as described previously (52). For immunoblot analysis, cells were washed twice with PBS before whole cell lysates were prepared by harvesting the cells with hot Laemmli buffer. Alternatively, cells were trypsin-shaved, harvested for mitochondrial preparation or subcellular fractionation, or subjected to neutral red uptake assay as further described. For VacA inhibitor experiments, AGS cells were treated with endocytosis inhibitors: cytochalasin D (20 µM, Sigma) or 4-bromobenzaldehyde N-(2,6-dimethylphenyl) semicarbazone (EGA; 20 µM, Sigma) (26); endosomal acidification inhibitors: bafilomycin A1 (62.5 nM, Invivogen), chloroquine (50 µM, Invivogen), ammonium chloride (5 mM, Sigma); proteasomal degradation inhibitor: MG132 (5 – 50 µM, Merck KgaA); cathepsin inhibitors: Pepstatin A methyl ester (10 or 20 µM, Merck, US1516485-1MG) or E64d (20 µM, Merck, 88321-09-9). All inhibitors were used both prior to (1 h) and during intoxication unless otherwise specified in figure legends. Culture media used in the aforementioned experiments was not supplemented with ammonium chloride unless otherwise stated.

### Culture and intoxication of human primary gastric organoids with VacA

This study was conducted in accordance with the Declaration of Helsinki, and the protocol was approved by the Cabrini Research Governance Office (CHREC 04-19-01-15) and the Monash Human Research Ethics Committee (MHREC ID 2518). Patient recruitment was led by surgeons in the Cabrini Monash University Department of Surgery. Gastric tissue was obtained from patients undergoing sleeve gastrectomy at the Cabrini Hospital, Malvern, Australia. All patients provided written consent.

Patient-derived normal human gastric tissue was placed in cold 1X chelating buffer (distilled water, 5.6 mM Na_2_HPO_4_, 8.0 mM KH_2_PO_4_, 96.2 mM NaCl, 1.6 mM KCl, 43.4 mM sucrose, 54.9 mM D-sorbitol), then cut in small fragments, and washed in 1X chelating buffer until the supernatant was clear. Glands were extracted by incubating tissue fragments for 10 mins at RT in 1X cold chelating buffer containing 10 mM EDTA, then resuspended in Matrigel (Corning, 356231) and seeded into 24-well tissue culture plates. Once polymerized, Matrigel was overlaid with 500 µL of culture medium composed of advanced Dulbecco’s modified Eagle medium (DMEM)/F12 (Gibco, Cat# 12634028) supplemented with penicillin/streptomycin (Gibco, Cat# 15140122), 10 mM HEPES (Gibco, Cat# 15630080), GlutaMAX (Gibco, Cat#35050061), 1xB27 (Gibco, Cat# 17504044), 100 ug/ml Primocin (Invivogen, Cat# ant-pm-2), 10 mM nicotinamide (Sigma, N0636), 1 mM N-Acetylcysteine (Sigma, A9165), 50 ng/mL recombinant human epidermal growth factor (EGF, PeproTech, Cat# AF-100-15), 10 % (v/v) noggin-conditioned medium, 10 % (v/v) R-spondin1-conditioned medium, 50 % (v/v) Wnt-conditioned medium, 200 ng/mL recombinant human fibroblast growth factor 10 (FGF10, PeproTech, Cat# 100-26-500), 1 nM gastrin (Tocris, Cat# G9145), 2 µM A-83-01 (Tocris, Cat# 2939). Following initial seeding of the cultures, 5 µM Y-27632 dihydrochloride kinase inhibitor (MedChemExpress, HY-10583), and 3 µM glycogen synthase kinase (GSK) 3β inhibitor (CHIR99021, Axon Medchem). Medium was refreshened every 2-3 days. Organoids were passaged by mechanical disruption every 6-10 days as described previously (53), or by enzymatic dissociation for seeding onto cell culture inserts as described below.

Transwell cell culture inserts (12 well, 16.38 mm height, 0.4 µm pore size) (Sarstedt, Cat# 83.3931.041) were placed in 12-well plates. Following the establishment of human gastric organoids in 24-well plates, organoids were mechanically scraped and collected into a tube with cold advanced DMEM/F12 (Gibco). Organoids were then centrifuged at 1500 rpm at 4° C for 5 mins, and supernatant discarded. Single cell suspension was obtained by treatment with TrypLE Express solution (Gibco, Cat# 12604013) for 3 minutes at 37° C. Cell pellet was resuspended in culture medium containing 5 µM Y-27632 dihydrochloride kinase inhibitor (MedChemExpress), and 150 µL of cell suspension per insert was seeded.

Cells were cultured in 150 µL apical and 1 mL basal complete medium for the first week or until a homogeneous monolayer of cells was formed, containing 5 µM Y-27632 dihydrochloride kinase inhibitor (MedChemExpress) for the first 3 days. Medium in the apical and basal compartments of the insert was refreshed every 2-3 days.

Human 2D gastric organoids were intoxicated with three-times concentrated *H. pylori* CS to a final concentration of 20 % (v/v) in both apical and basal media compartments at 37 ^°^C for 24 h. For immunoblot analysis, 2D organoids were gently washed with PBS before whole cell lysates were prepared by harvesting 2D organoids in transwells with hot Laemmli buffer.

### Immunoblot analysis and antibodies

Samples denatured by boiling in Laemmli buffer containing 100 mM DTT were separated in 12 % SDS-PAGE gels according to the Laemmli buffer system and electrophoretically transferred to nitrocellulose membranes for immunoblot analysis. Primary antibodies used were rabbit anti-p33 and anti-p55 (this study); rabbit anti-F1β (54) and anti-Sam50 (55); rabbit anti-Strep tag II and mouse anti-rab5 (Abcam). Secondary antibodies were ECL anti-rabbit or anti-mouse IgG HRP-conjugated (Amersham), and bound secondary antibody was detected using ECL Prime chemiluminescent substrate (Amersham).

### Isolation of AGS endosomal/mitochondrial-enriched fraction

AGS cells were washed twice with ice-cold PBS and all subsequent steps were performed on ice or at 4 °C. Cells harvested by scraping were pelleted by centrifugation at 1000 x*g* for 10 min and resuspended in isotonic Solution A (20 mM HEPES pH 7.5, 220 mM mannitol, 70 mM sucrose, 1 mM EDTA and freshly added 0.2 mg/mL PMSF). Cells were lysed by repeated expulsion through a 22-gauge needle. Cell lysates were clarified by centrifugation at 1000 x*g* for 15 min and the resulting supernatant then centrifuged at 10,000 x*g* for 30 min to pellet mitochondria. The mitochondrial pellet was resuspended in Solution A and centrifuged again at 1000 x*g* for 15 min at 4 °C to pellet any remaining whole cells and contaminating nuclei. Mitochondria-enriched sample protein concentrations were determined by BCA assay (Pierce) and stored at −80 °C.

### Mass spectrometry

SDS-polyacrylamide gel blocks corresponding to VacA fragments identified by parallel VacA immunoblot analysis were excised and subjected to in-gel digestion using trypsin (Promega) or chymotrypsin (Promega) and LC-MS/MS analysis as previously described (56). Mass spectrometric raw data was processed and searched by Byonic software (Protein Metrics, v2.6.46) (57) or MASCOT (v2.4). Search parameters include fully specific or semi specific tryptic/chymotryptic cleavage sites with up to 3 missed cleavages, cysteine carbamidomethylation as fixed modification and methionine oxidation as common variable modification. Database searching was performed against *H. pylori* 26695 protein sequences downloaded from Uniprot (Sep 2015). Mass spectrometry peptide scores, data from Western blot and mutant analyses together with the protein molecular weight calculator, Compute pI/Mw (58) were used as criteria to predict the N- and C-terminal boundaries of the VacA fragments.

### Subcellular fractionation

VacA^+^- or VacA^−^-treated AGS cells were scraped from a 175 cm^3^ flask into 0.5 mL of ice-cold homogenization buffer containing 250 mM sucrose, 1 mM EDTA, 10 mM Tris-HCl pH 7.4, mini c0mplete™ EDTA-free protease inhibitor cocktail (Roche). Cells were homogenized on ice by repeated passage through a 22-gauge needle and centrifuged at 800 x*g* at 4 °C for 10 min to pellet whole cells and nuclei. The resulting supernatant was overlaid on a 5-step discontinuous sucrose gradient (2.28 mL each of 18%, 27%, 35%, 43% and 50% (w/v) sucrose solutions in 10 mM Tris-HCl pH 7.4, 1 mM EDTA) and centrifuged at 100,000 x*g* at 4 °C for 19 h using a SW 40 Ti rotor. Gradients were collected from the top of the gradient into 12 equal fractions using a Brandel density gradient fractionation system (Brandel/ISCO). The protein component of each fraction was precipitated by 10 % (v/v) TCA followed by acetone washes, separated by SDS-PAGE, and analyzed by immunoblot with various antibodies as specified in figure legends.

### Trypsin-shaving of AGS cells

AGS cells in 24-well plates were washed twice with 500 µL/well of pre-warmed PBS before incubation with 150 µL/well of pre-warmed trypsin solution (0.25% (w/v) trypsin, 0.91 mM EDTA, Gibco) for 5 mins at 37 °C in the presence of 5 % CO_2_. Trypsin activity was quenched by the addition of 450 µL/well of pre-warmed DMEM supplemented with 10 % HI-FBS. AGS cells were centrifuged at 150 x*g* and the pellet resuspended in 1 mL of ice-cold PBS. The previous step was repeated once more before the cell pellet was homogenized in hot SDS Laemmli buffer and analyzed by immunoblot.

### Quantification of vacuolation

Neutral red dye uptake (NRU) assays were performed in 96-well plates as previously described (59, 60) with modifications. Briefly, 0.5 % neutral red dye (Sigma-Aldrich) stock solution was diluted in phenol red-free RPMI supplemented with 10 % HI-FBS and 20 mM HEPES pH 7 to make a 0.025 % neutral red dye working solution. AGS cells were treated with VacA^+^, VacA^−^ or cell-culture medium (mock-treated) in triplicate wells per treatment condition for varying durations, as specified in figure legends. For pulse-chase NRU experiments in which the cells were pulsed with VacA^+^ for 30 minutes, the culture media was supplemented with 5 mM ammonium chloride for the final 30 mins of incubation to potentiate vacuolation. For both accumulation and pulse-chase experiments, treatments were staggered so that all time-points finished simultaneously and all wells contained cells of the same culture duration post-seeding. At completion of treatment, cells were washed once with 150 µL/well pre-warmed PBS and incubated with 100 µL/well of pre-warmed neutral red dye working solution for 4 min at room temperature. Excess dye was removed by washing cells twice with 150 µL/well pre-warmed PBS and the dye extracted with 100 µL/well of acidified alcohol (50 % (v/v) ethanol in 1 % (v/v) acetic acid). Within each experimental cohort, optical densities were read at 540 nm and the values standardized by subtraction of the median value of six parallelly mock-treated wells of AGS cells. Each experiment was performed three times with triplicate wells per condition included in each independent experiment.

### 2D-PAGE of AGS endosomal/mitochondrial-enriched fraction

Mitochondria-enriched fraction isolated from VacA^+^-treated AGS cells (24 h treatment) was resuspended in digitonin lysis buffer (1 % (v/v) digitonin, 20 mM Tris HCl pH 7.4, 0.1 mM EDTA, 50 mM NaCl, 10 % (v/v) glycerol, 1 mM PMSF) and incubated on ice for 30 min with periodic vortexing. Samples were then centrifuged at 10,000 x*g* for 30 min and the solubilized mitochondrial proteins were collected in the supernatant. For the first dimension electrophoresis, the supernatant was combined with blue native (BN) sample buffer (final 1X BN buffer of 1 % (w/v) Coomassie Brilliant Blue G-250, 100 mM 6-aminocaproic acid, 20 mM Bis-Tris pH 7) and electrophoresed on a 4 – 16.5 % BN polyacrylamide gel. For the second dimension, the BN lane containing mitochondrial proteins was excised and washed gently for 1 min in SDS-PAGE running buffer, then encased orthogonally in a 4 % SDS-polyacrylamide stacking gel overlaying a 12 % SDS-polyacrylamide gel. SDS-PAGE and immunoblot analysis were performed as described.

### Statistical analysis

All statistical analysis was performed using Prism v 8.4.3 (GraphPad) and p-values <0.05 were considered significant. Neutral red uptake data was statistically assessed in response to each treatment over time by comparison with t = 0, wherein elevated optical density indicated vacuolation and reduced optical density indicated reduced cell viability. For the accumulation experiments (*SI Appendix*, Figure S6c), the median data from each of three independent experiments was stacked in a spreadsheet sub-column and assessed by repeated measures two-way ANOVA with the Geisser-Greenhouse correction. For the pulse-chase NRU experiments (*SI Appendix*, Figure S7a), the 0.5 – 10 h time-points and 16 – 24 h time-points were assayed separately so the data was left unmatched and analyzed by ordinary two-way ANOVA. In each NRU analysis, the two intra-assay covariates of treatment and time were assessed using Holm-Sidak multiple comparisons test with individual variances computed for each comparison, and the residuals were confirmed to distribute horizontally in residual and homoscedasticity plots, and diagonally in a QQ plot.

Non-parametric analysis of a correlation between VacA fragments and vacuolation was performed by determining the Spearman r of VacA anti-p55 densitometry values against the NRU values obtained at identical time-points. Data in this analysis was compiled from 3 independent experiments: NRU data (x-direction) was a single median value per time (independent replicates shown in *SI Appendix*, Figure S6a); densitometry data (y-direction) was each independent experimental value per time point. The 0.5 – 10 h and 16 - 24 h densitometry data were analyzed separately because they originated from different sets of immunoblots, but only the 0 – 10 h data had enough datapoints to generate p-values. Cubic spline curves generated with three knots were included in the correlation plots to aid in visualizing the correlation but are not part of the analysis.

## Availability of data and biological materials

The data that support the findings of this study and the biological materials unique to this study are available from the corresponding author upon reasonable request.

## Acknowledgments

We thank the Monash Animal Research Platform for the production of the VacA-reactive antisera, and Dr David Steer and the Monash Proteomics and Metabolomics Facility for their assistance with MS analyses. We thank the Monash Organoid Program for their assistance with human gastric organoid culture, and Dr Thomas Naderer for providing anti-Rab5. The project was supported by funding from the National Health and Medical Research Council (NHMRC Project 2010001749; NHMRC Program 1016629), the National Institutes of Health (AI039657), and the Department of Veterans Affairs (BX004447).

## Supplementary Information

**Figure S1.**
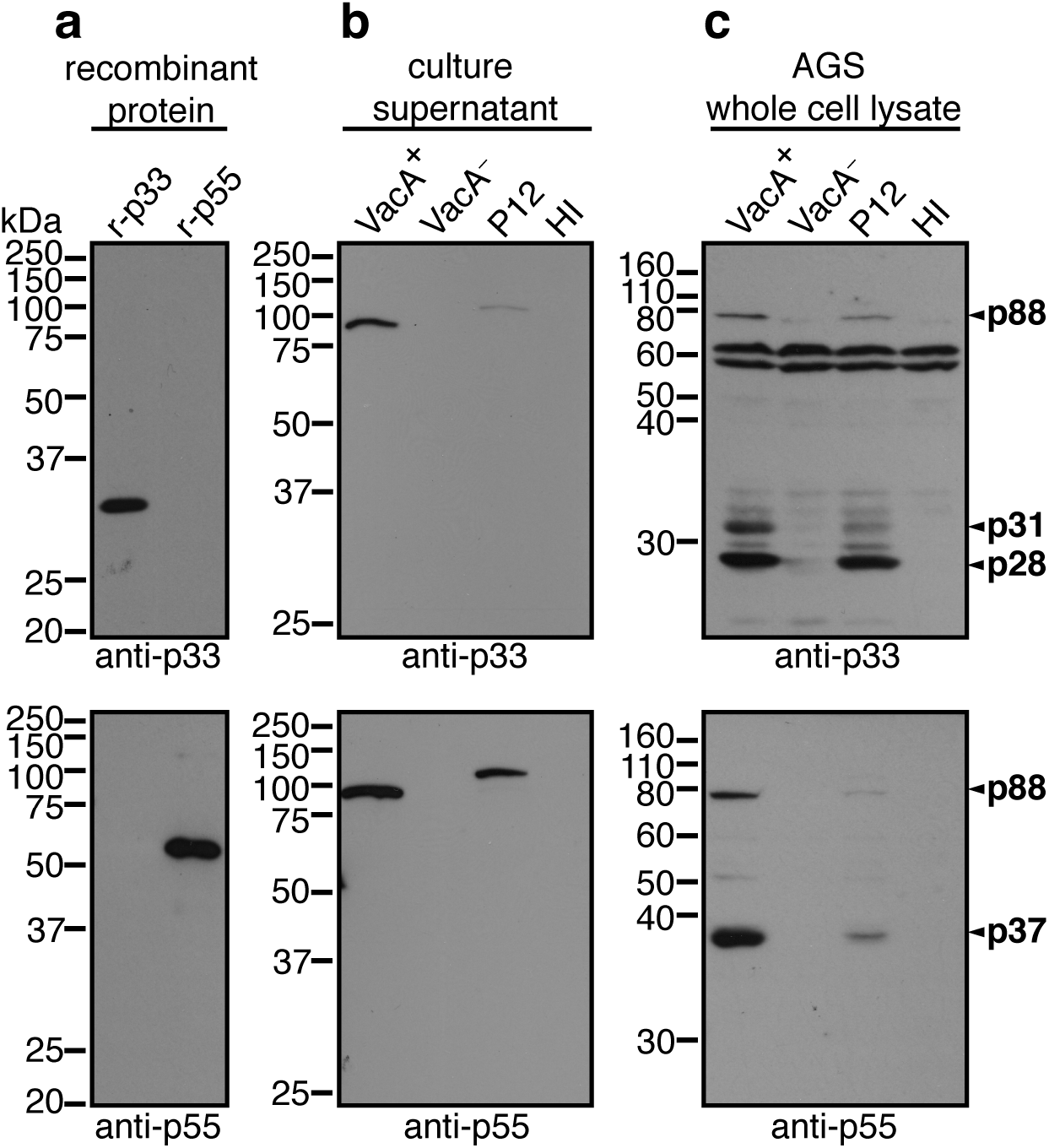
VacA antisera are reactive with recombinant p33 and p55, native secreted VacA and novel intracellular cleavage products. **a**, Recombinant purified p33 (r-p33) and p55 (r-p55) proteins, **b**, *H. pylori* CS from VacA^+^*Hp* (VacA^+^), VacA^−^*Hp* (VacA^−^), P12 or HI broth, and **c**, AGS whole-cell lysates post-24 h treatment with the indicated CS or HI broth were subjected to immunoblot analysis using anti-p33 or anti-p55 as indicated.

**Figure S2.**
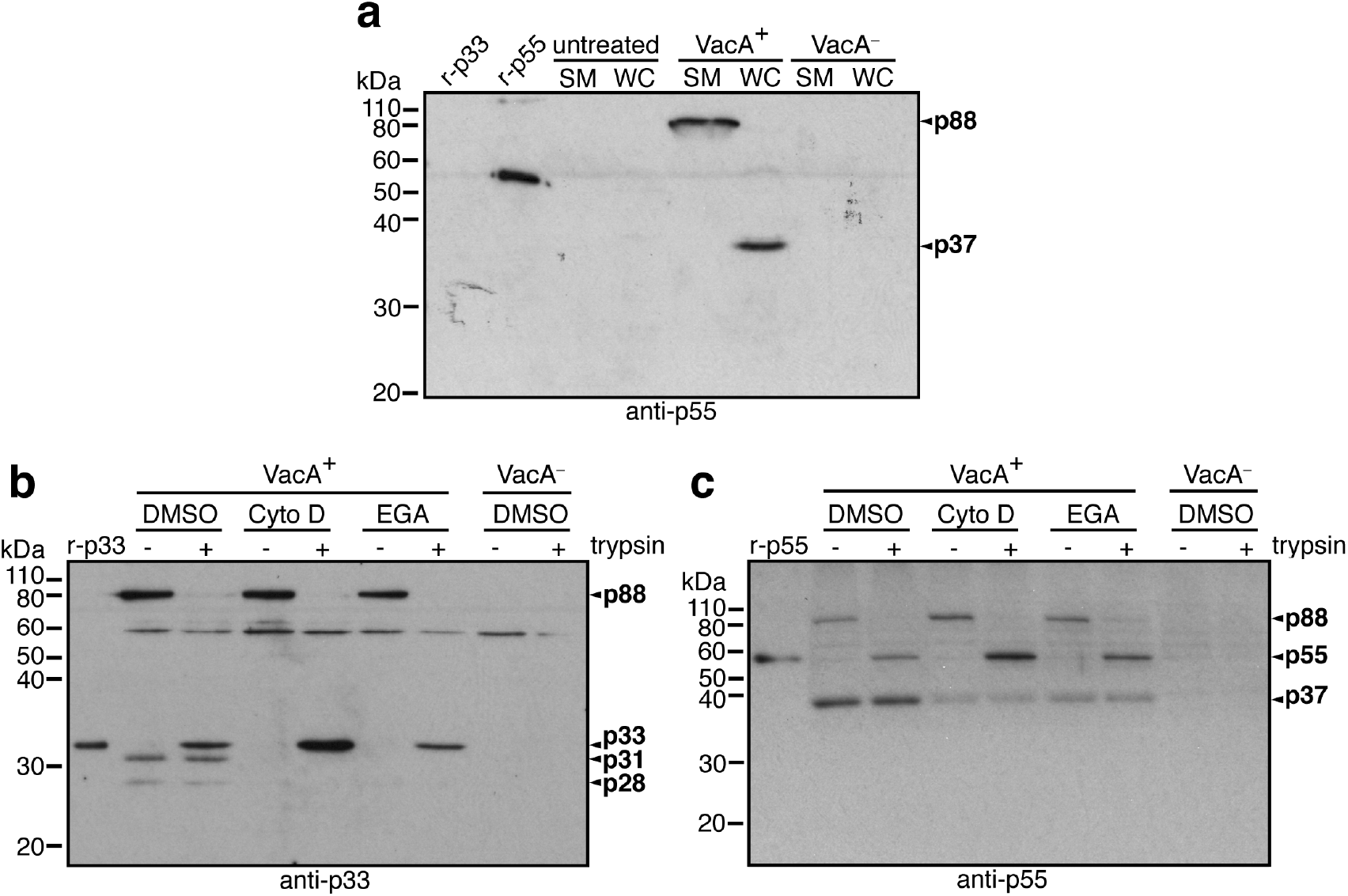
VacA fragments are generated inside host cells. **a**, Immunoblot analysis of AGS cells treated with VacA^+^ or VacA^−^ for 24 h before analysis of AGS spent culture medium (SM) and whole cell lysate (WC) using anti-p55. No p37 was detected in SM suggesting that VacA fragments do not arise extracellularly. Immunoblot analysis using **b**, anti-p33 or **c**, anti-p55 of VacA^+^-treated (6 h) AGS cells without (DMSO) or with endocytosis inhibitor (20 µM cytochalasin D or 20 µM EGA). Cells were harvested without (-) or with (+) trypsin to cleave extracellular VacA p88 into p33 and p55 as a means to distinguish between extracellular and intracellular p88. The susceptibility of p88 to trypsin confirms that VacA internalization was blocked by these inhibitors, and the absence of processed VacA fragments p31, p28 and p37 in cells treated with cytochalasin D or EGA further suggests that VacA is processed intracellularly.

**Figure S3.**
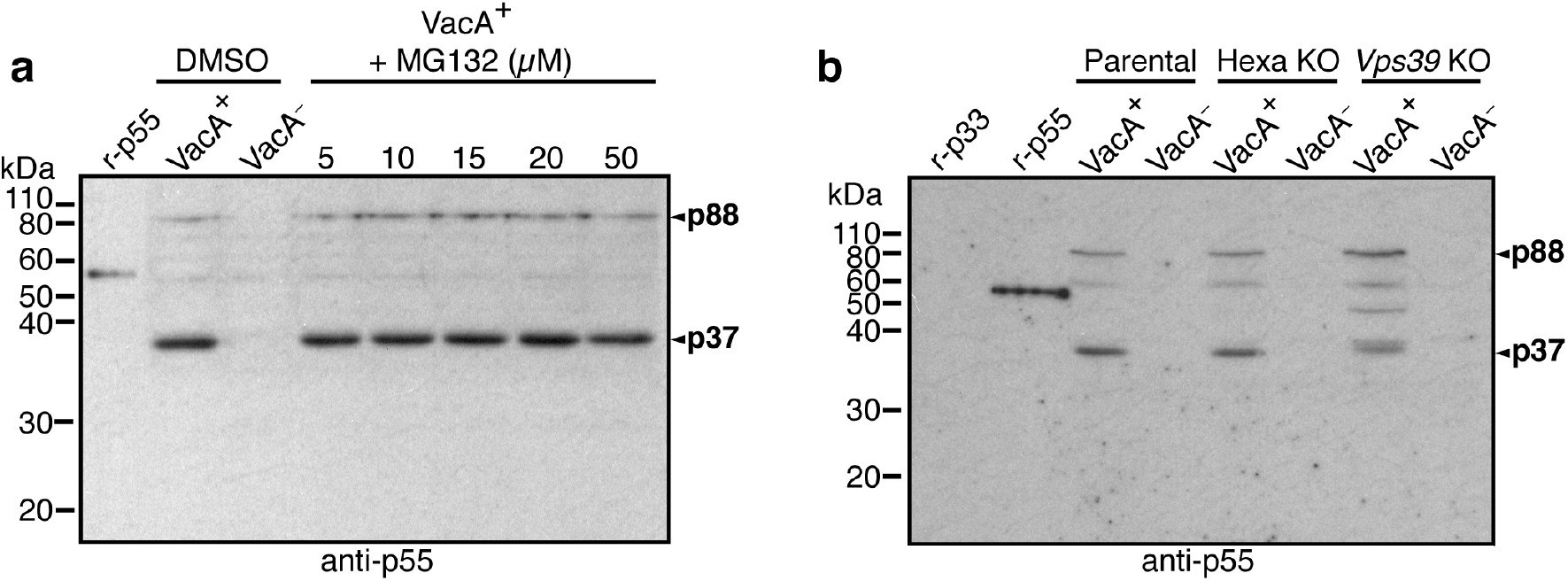
Endo-lysosomal fusion, but not autophagy or the proteasome, is involved in intracellular VacA processing. **a**, Immunoblot analysis of AGS cells pre-treated with various concentrations of MG132 as indicated for 1 h before incubation with VacA^+^ for 6 h; blot probed using anti-p55. Processing of VacA p88 into p37 was unaffected by MG132 at all concentrations tested suggesting that proteasomal degradation is not involved in VacA processing. **b**, Immunoblot analysis of HeLa parental cells, Hexa KO and Vps39 KO cells incubated with VacA^+^ or VacA^−^ for 16 hours using anti-p55. Processing of VacA p88 was unaffected in Hexa KO cells suggesting that autophagy is not involved in VacA processing. However, in cells lacking Vps39, a key gene involved in late endosomal and lysosomal clustering and fusion, VacA processing was affected as shown by the presence of an intermediate band at ∼45 kDa, doublet bands of ∼38/39 kDa and lower overall p37 levels. These data suggest that endosomal maturation is important for complete cleavage of VacA.

**Figure S4.**
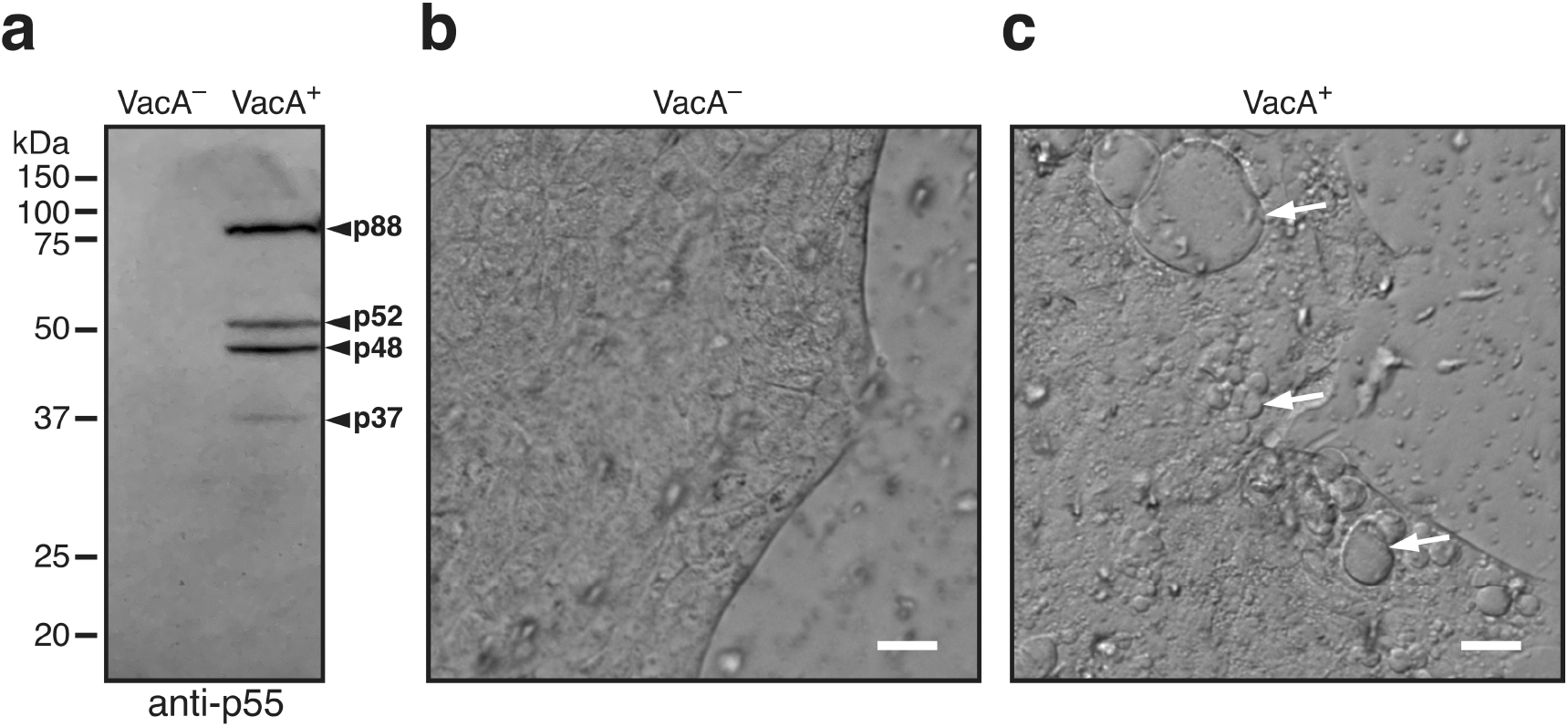
VacA is processed by and induces vacuolation in human 2D primary gastric organoids. **a**, Immunoblot analysis of VacA^−^ and VacA^+^-treated human 2D primary gastric organoid monolayers using anti-p55; VacA p88 is processed by human 2D primary gastric organoids into smaller fragments (fragments p52 and p48 likely represent intermediate products). Transmitted-light micrographs of **b**, VacA^−^ or **c**, VacA^+^-treated human 2D primary gastric organoid monolayers; Vacuoles indicated by arrows; data shown are representative of three independent experiments (Scale bar, 20 µm).

**Figure S5.**
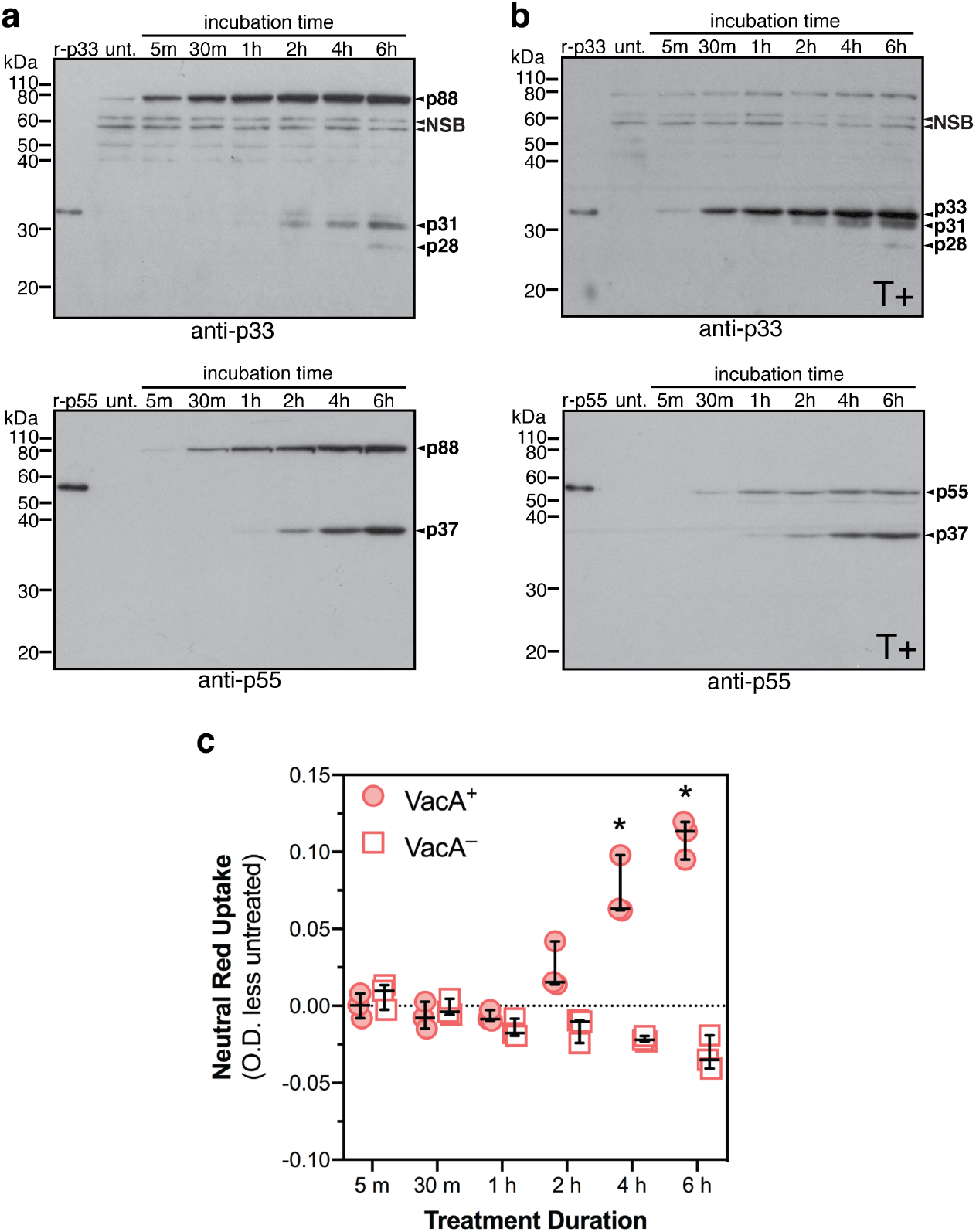
Generation and accumulation of VacA fragments in gastric epithelial cells correlates with vacuolation – accumulation experiments. Immunoblot analysis using anti-p33 or anti-p55 (as indicated) of AGS cells treated with VacA^+^ for various times before harvest **a**, without trypsin-shaving or **b**, with trypsin-shaving (T+). **c**, Neutral red uptake (NRU) assay of AGS cells treated with VacA^+^ or VacA^−^ for various times; y-axis denotes culture supernatant-treated cells O.D._540 nm_ minus untreated control O.D._540 nm_; each data-point is median of biological triplicates from a single experiment; bars denote median ± 95% CI of three independent experiments; statistical comparison of each dataset against starting NRU levels (t = 5 minutes), repeated measures 2-way ANOVA *P< 0.05; p-values of all comparisons provided in *SI Appendix*, Table S2. Generation and accumulation of VacA fragments coincides with and is directly proportional to the development of vacuolation, whereas p88 is vulnerable to trypsin at all time points as shown by the presence of p33 and p55 products in cells harvested with trypsin. These data suggest that p88 accumulates on the plasma membrane, is processed into fragments upon internalization and that these fragments represent the intracellular functional form of VacA.

**Figure S6.**
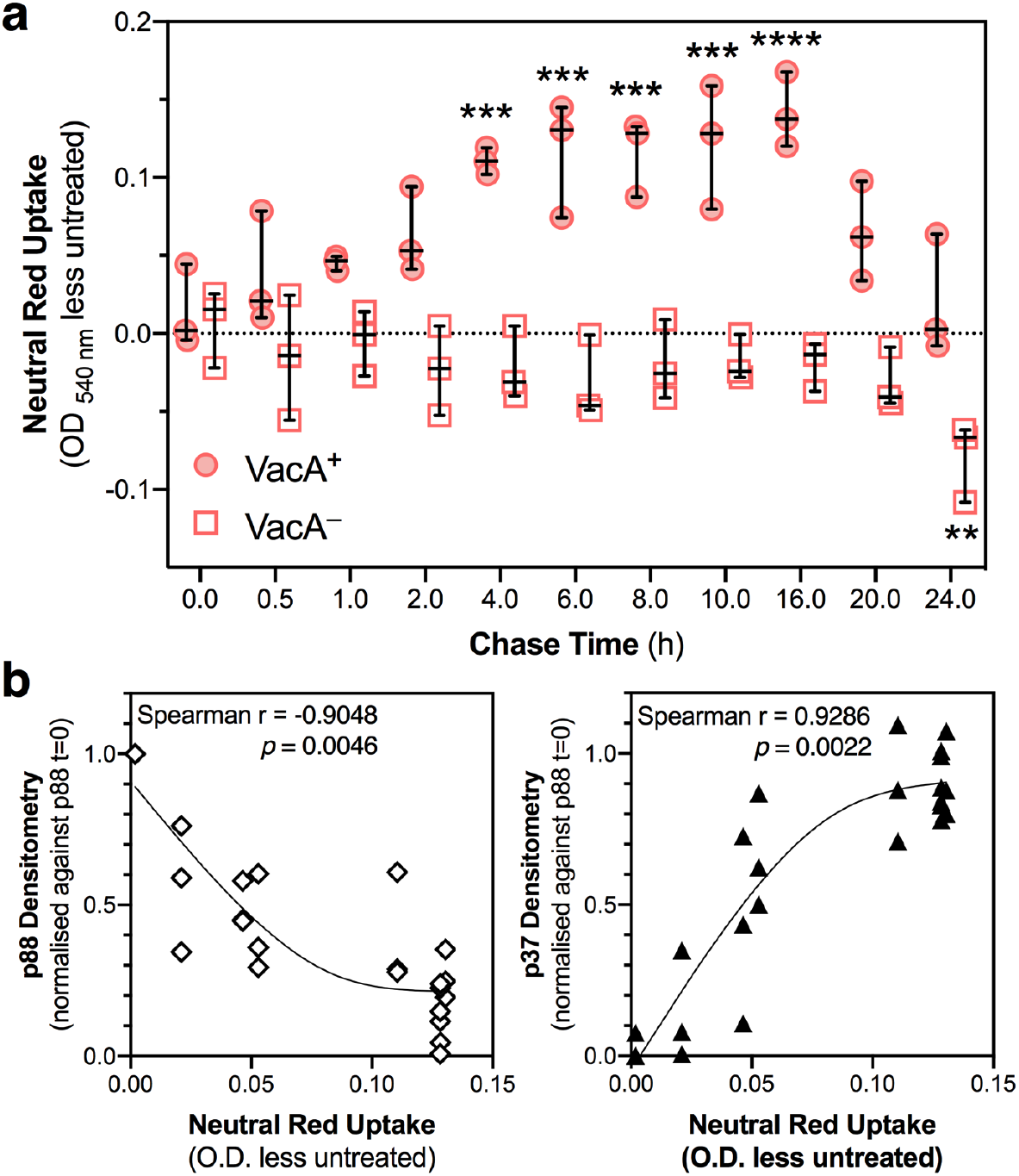
Initiation and resolution of vacuolation correlates with VacA fragment levels in gastric epithelial cells – pulse-chase experiments. **a**, NRU assay of AGS cells pulsed with VacA^+^ or VacA^−^ for 30 min at staggered time-points and chased (after washing) with fresh media for various times from 0 to 24 h post-pulse before synchronized NRU assay. Vacuolation was potentiated by supplementing media with 5 mM NH_4_Cl 30 min prior to NRU assay. Significant vacuolation was evident from 4 h to 16 h post-chase with VacA^+^, but not with VacA^−^ at any time-point. By 20 h post-chase, NH_4_Cl was no longer able to significantly potentiate vacuolation in response to VacA^+^ intoxication. y-axis denotes CS-treated cells O.D._540 nm_ minus untreated control O.D._540 nm_; each data-point is median of biological triplicates from a single experiment; bars denote median ± 95% CI of three independent experiments; statistical comparison of each dataset against base-line NRU levels (t = 0 h), ordinary 2-way ANOVA **p < 0.01, *** p <0.001, **** p <0.0001; actual p-values of all comparisons provided in *SI Appendix*, Table S3. **b**, Vacuolation development negatively correlated with the level of cell-associated p88, but positively correlated with the level of intracellular processed VacA (p37). Spearman correlation analysis was performed using median NRU data and individual densitometry data-points, all from 3 independent experiments and 0 – 10 h post-chase.

**Figure S7.**
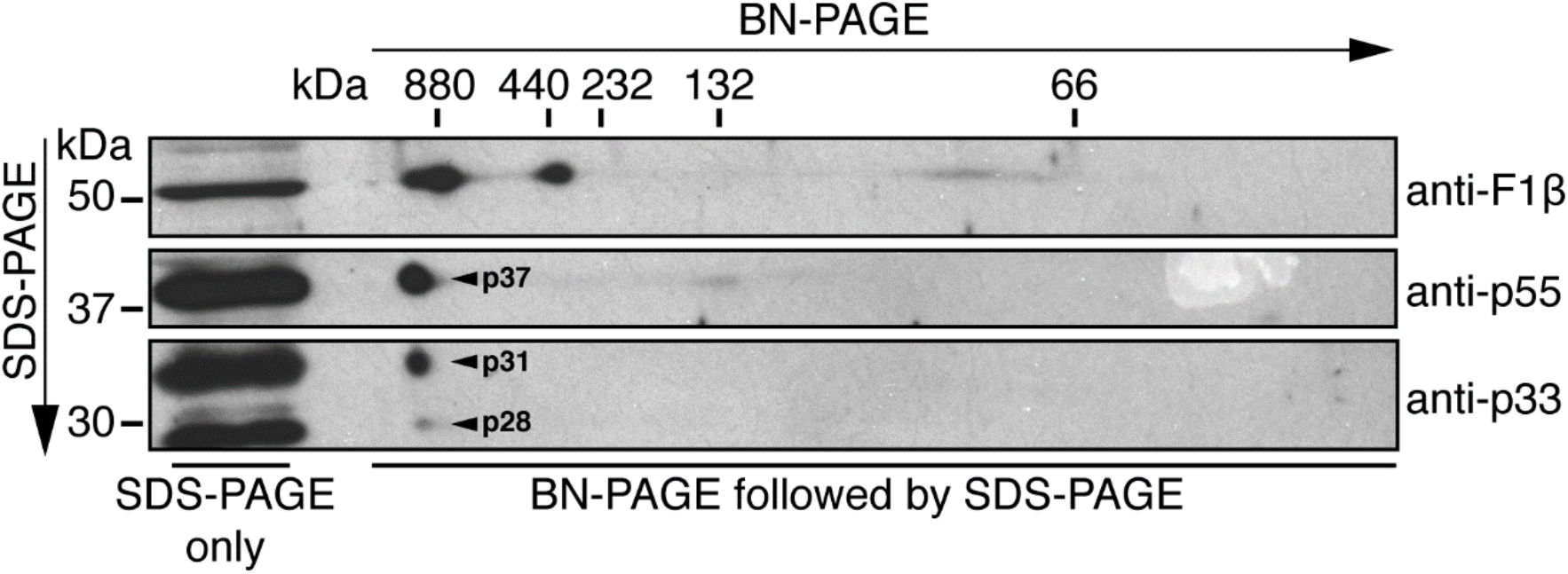
p31, p28 and p37 are constituents of higher-order complexes in mitochondria-enriched AGS fractions. Independent repeat of experiment shown in Fig. 4a: 2D-PAGE and subsequent immunoblot analysis of mitochondria-enriched fractions isolated from AGS cells treated with VacA^+^ for 24 h using the indicated antibodies. Processed products p31, p28 and p37 are constituents of the same higher-order VacA complex that may represent functional VacA anion channels; F_1_β is the beta subunit of F_1_-ATPase complex and serves as a BN-PAGE control.

**Figure S8.**
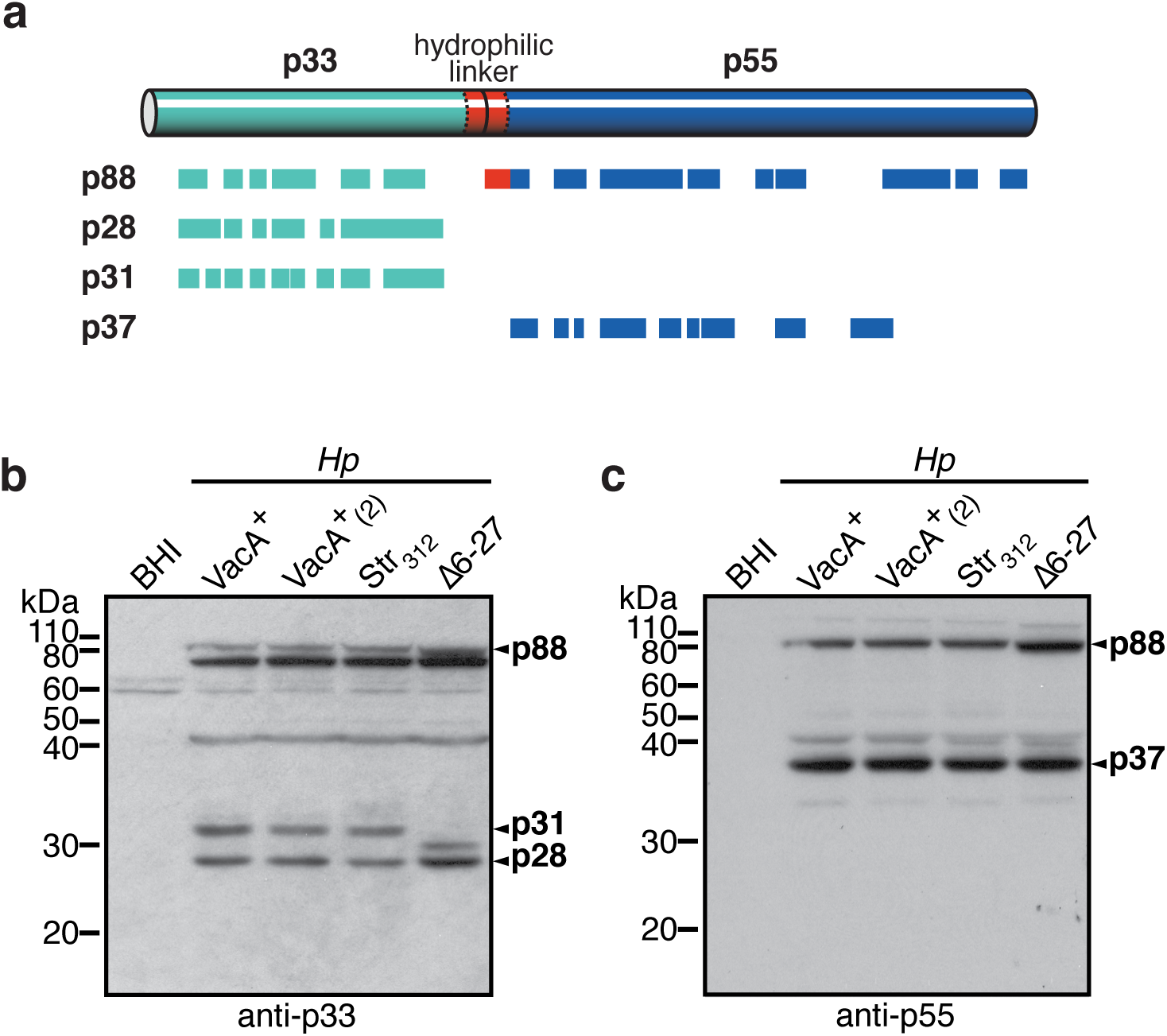
Identification of novel VacA cleavage products and difference between p31 and p28. **a**, VacA coverage map of tryptic/chymotryptic peptides identified following common bottom-up MS analyses of p88, p31, p28 or p37 in-gel bands. The origins of these fragments were further confirmed by epitope-tagging the predicted p31 and p37 regions of VacA, which conferred anti-Strep reactivity to either the anti-p33 or anti-p55 reactive bands (*SI Appendix*, Fig. S10). Immunoblot analysis of AGS cells infected with various *H. pylori* 60190-derived strains for 24 h using **b**, anti-p33 or **c**, anti-p55; VacA^+^(2) designates alternative 60190wt strain, Str_312_ contains a Strep-tag within the hydrophilic linker at aa position 312 and Δ6-27 lacks aa residues 6-27. Similar to VacA^+^, p88 from strains VacA^+^(2) and Str312 were processed into fragments p31, p28 and p37 as expected (panels b,c). Interestingly, the presence of a smaller product in place of p31 in strain Δ6-27 (panel b), which lacks the hydrophobic N-terminal alpha helix, suggests that the difference between fragments p31 and p28 corresponds to the presence and absence, respectively, of this region.

**Figure S9.**
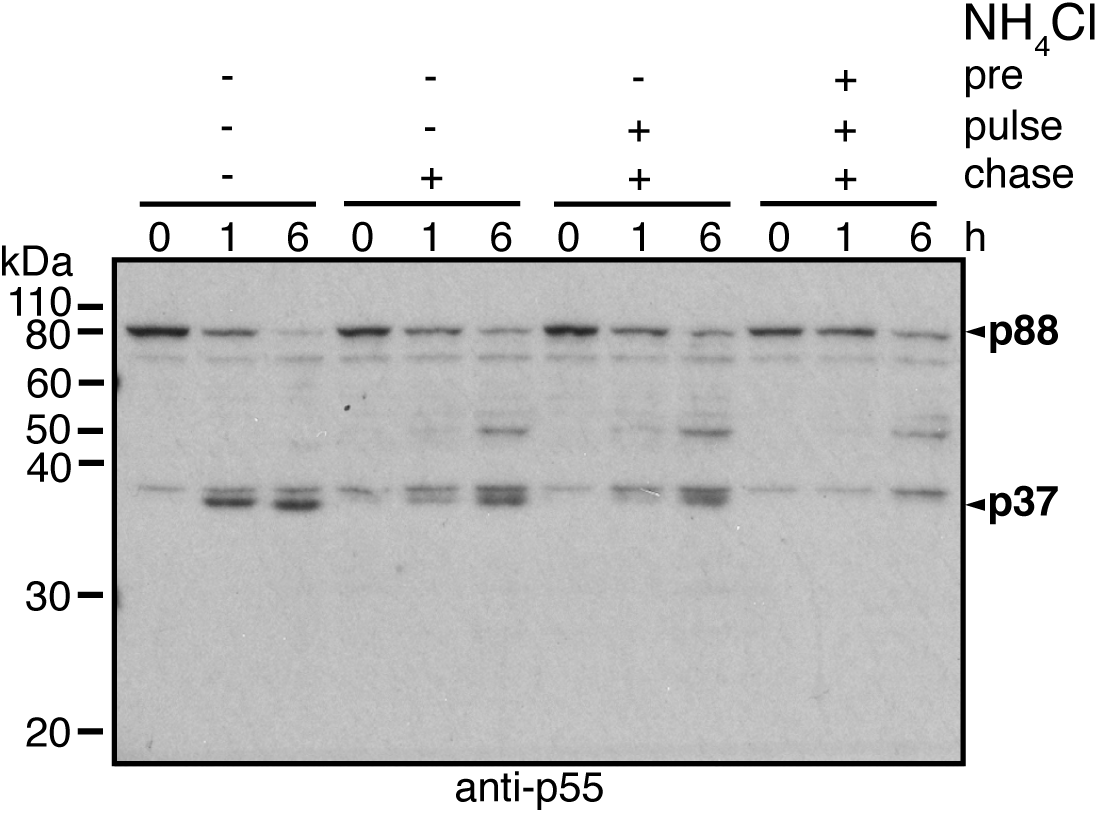
NH_4_Cl reduces VacA processing. Immunoblot analysis of VacA^+^ pulse-chase time-course in AGS cells without (-) or with (+) 5mM NH_4_Cl during different stages of the pulse-chase: pre (1 h pre-treatment), pulse (0.5 h co-pulse with VacA^+^) or chase (6 h). Samples were harvested at 0, 1 or 6 h post-pulse as indicated. In the absence of NH_4_Cl p88 was processed into p37 as soon as 1 h into the chase and accumulated over time. Conversely, almost no p88 remained after 6 h. When NH_4_Cl was added during the chase alone or during the pulse and chase stages, less processing of p88 occurred resulting in lower p37 levels and higher p88 levels after 6 h. Interestingly, under these conditions an intermediate product of ∼ 50 kDa was detected after 6 hours suggesting that VacA likely undergoes sequential processing. When cells were treated with NH_4_Cl during the pre-treatment, pulse and chase stages, processing of p88 into p37 was completely inhibited and p88 levels remained high throughout the chase. Again, an intermediate ∼ 50 kDa product was detected after 6 h, which suggests that NH_4_Cl does not completely inhibit VacA processing. These data show that NH_4_Cl inhibits host cell VacA processing to varying degrees depending on when the drug is added and also demonstrates that VacA processing occurs in a sequential, step-wise manner.

**Figure S10.**
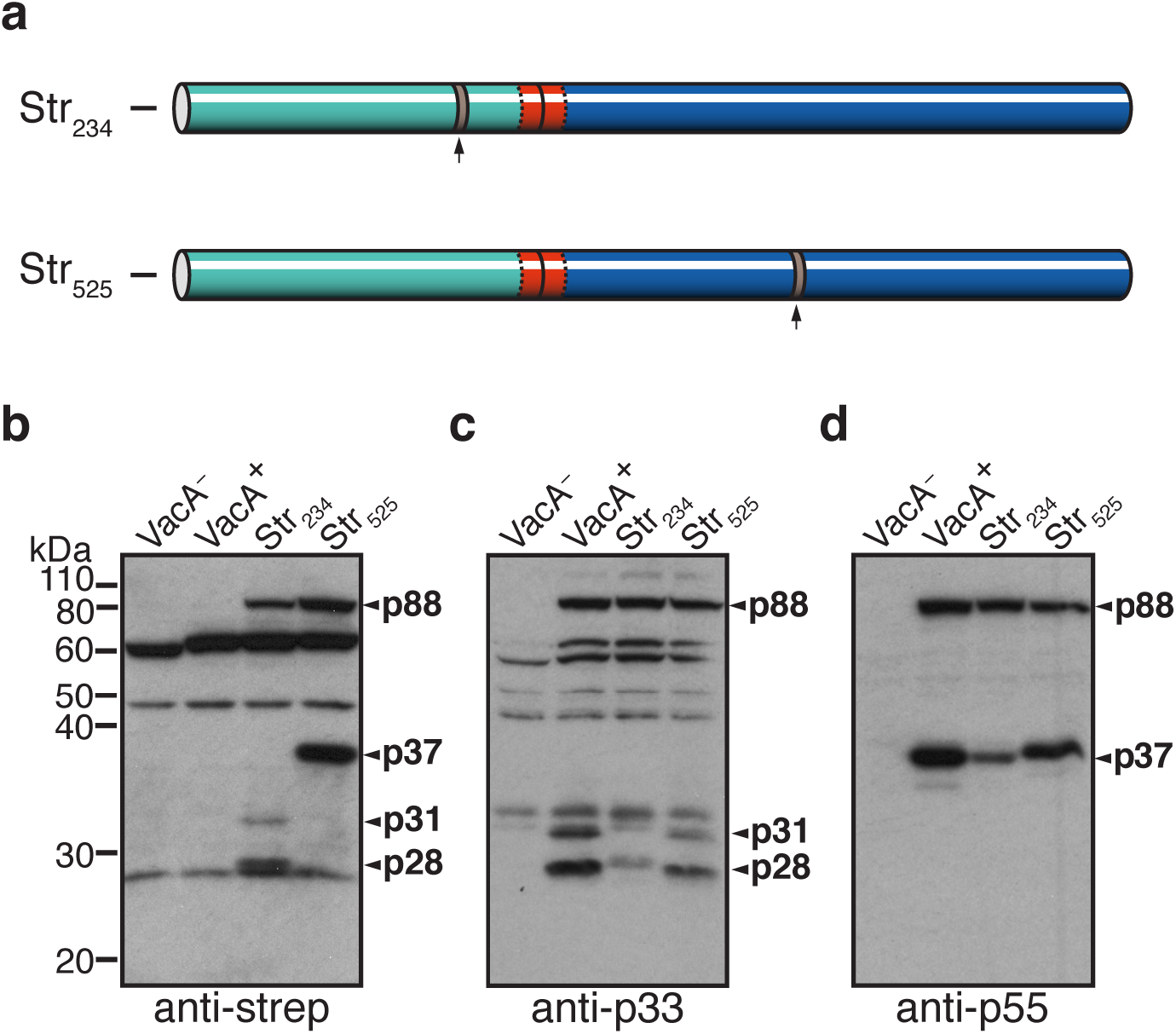
Insertion of Strep-tag into p31/p28 or p37 VacA regions further confirms their VacA origin. **a**, Strep-tags were inserted into 60190 VacA at aa positions 234 (Str_234_) or 525 (Str_525_) to further confirm the VacA origin of p31/p28 and p37 fragments, respectively; arrow indicates position of Strep-tag. Immunoblot analysis of AGS cells treated for 24 h with VacA^−^, VacA^+^, Str_234_ or Str_525_ CS using **b**, anti-Strep **c**, anti-p33 or **d**, anti-p55. VacA p31 and p28 products were detected using anti-Strep following treatment of host cells with Str_234_, while p37 was detected using anti-Strep following treatment of host cells with Str_525_ (panel b). Furthermore, treating cells with Str_234_ resulted in slightly larger p31 and p28 fragments compared to corresponding VacA^+^ fragments when probed with anti-p33 (panel c), while cells treated with Str_525_ resulted in a larger p37 fragment when probed with anti-p55 (panel d). Together these data demonstrate the successful insertion of Strep-tags into p31, p28 and p37 fragments, further confirming their VacA origin.

**Table S1.**
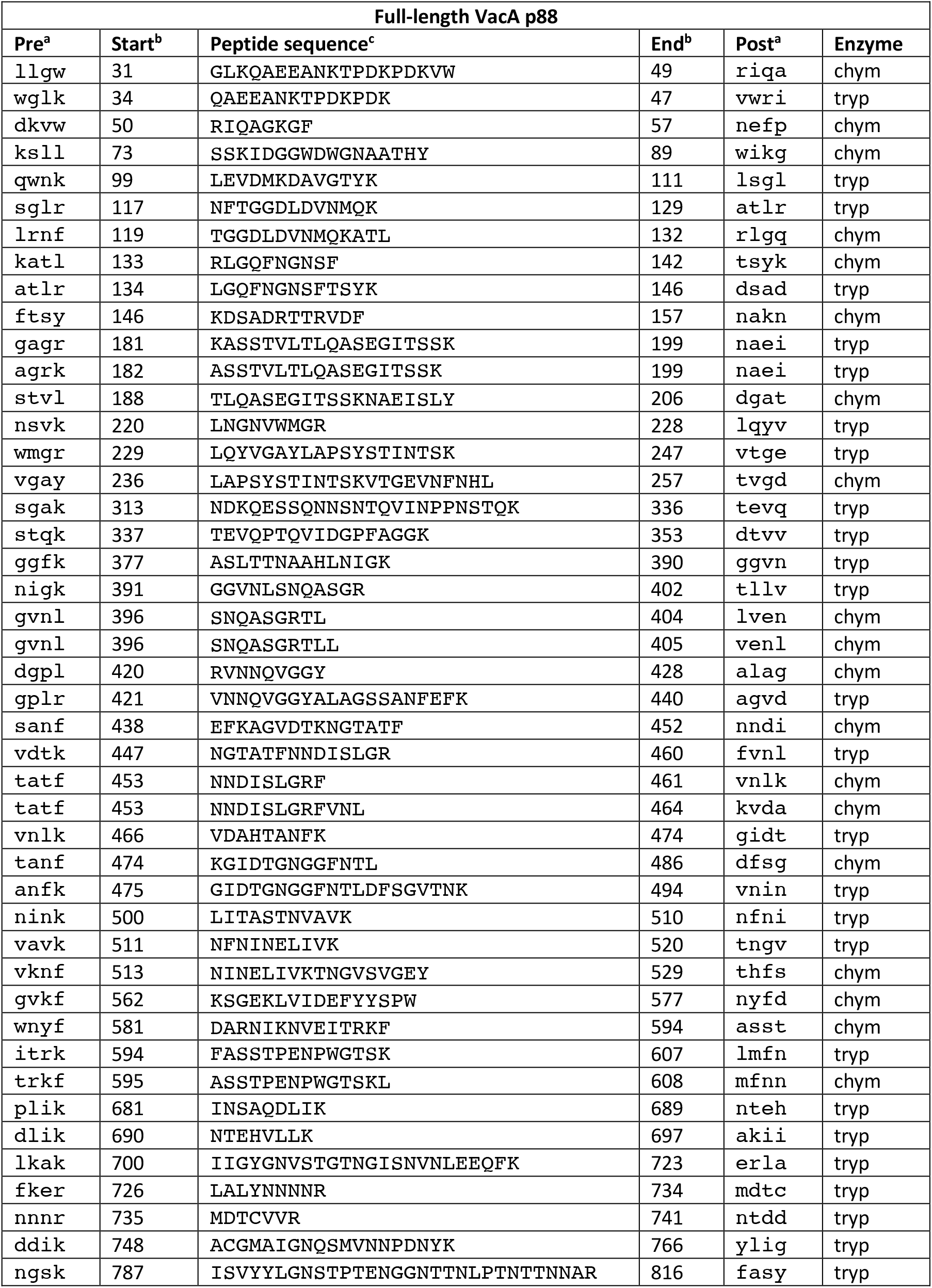
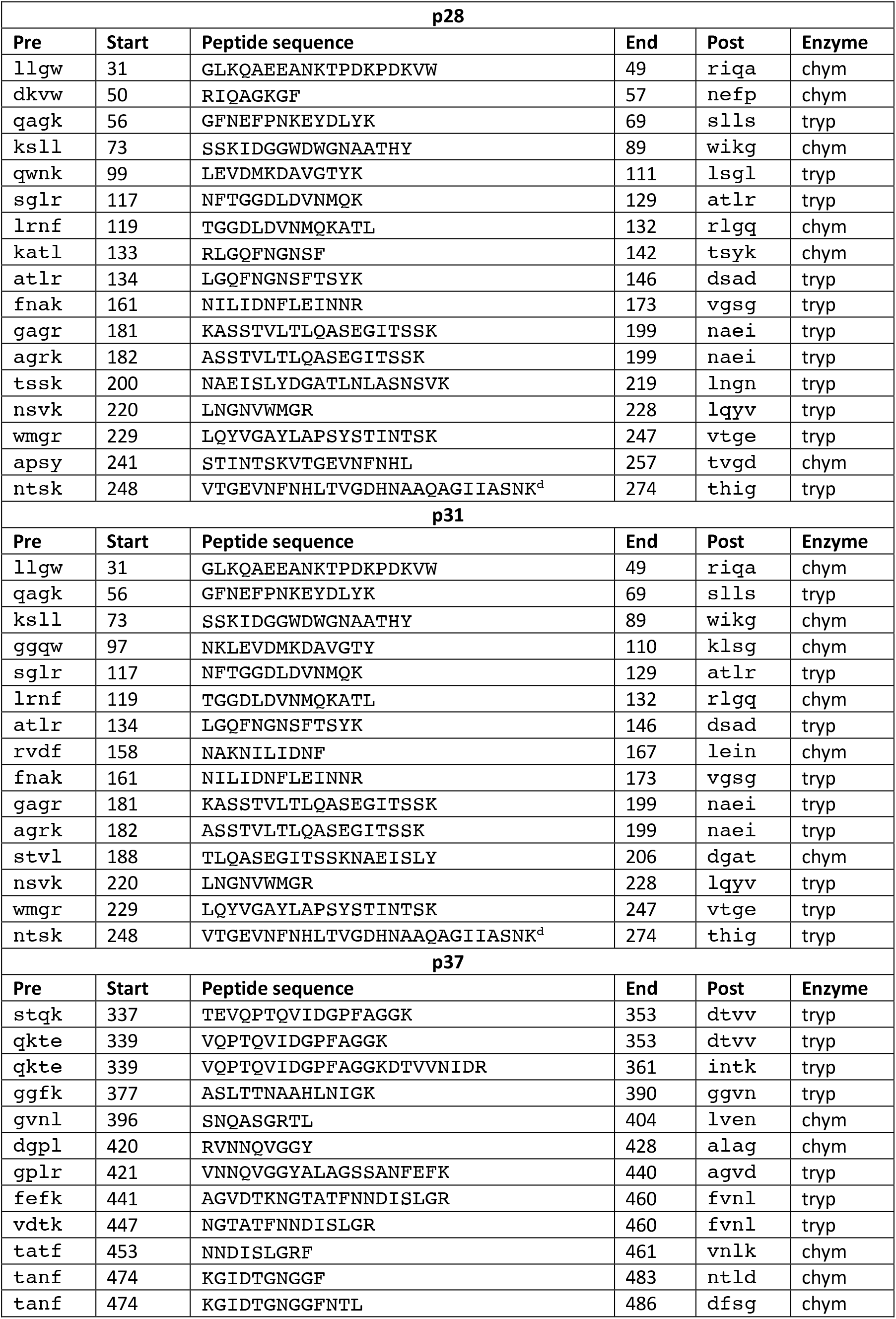
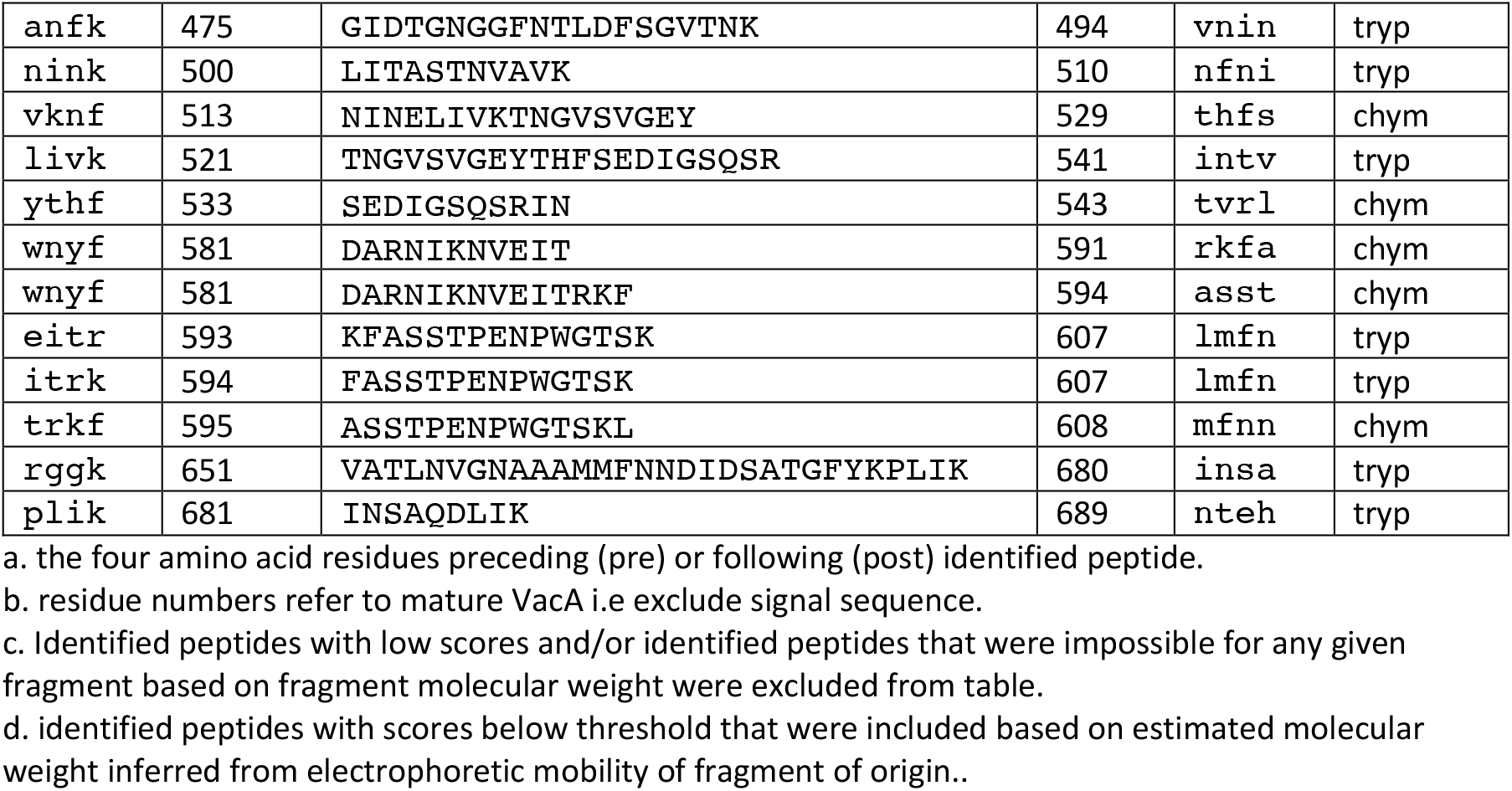
Tryptic (tryp) or chymotryptic (chym) peptides detected following common bottom-up MS analyses of in-gel VacA bands

**Table S2.** Accumulation neutral red assay – individual adjusted p-values (significant p-values bolded).

| Treatment | Comparison of 5 minutes versus all later times |  |  |  |  |
| --- | --- | --- | --- | --- | --- |
|  | times post-intoxication (adjusted p-values) |  |  |  |  |
|  | 30 m | 1 h | 2 h | 4 h | 6 h |
| VacA<sup>+</sup> | 0.2962 | 0.2962 | 0.1428 | **0.0466** | **0.0252** |
| VacA<sup>-</sup> | 0.3963 | 0.0888 | 0.1984 | 0.1148 | 0.0888 |

**Table S3.** Pulse-Chase neutral red assay – individual adjusted p-values (significant p-values bolded).

| Treatment | Comparison of 0 minutes versus all later times post-pulse<br>(adjusted p-values) |  |  |  |  |  |  |  |  |  |
| --- | --- | --- | --- | --- | --- | --- | --- | --- | --- | --- |
|  | 0.5 | 1 | 2 | 4 | 6 | 8 | 10 | 16 | 20 | 24 |
| VacA<sup>+</sup> | 0.54 | 0.43 | 0.14 | **0.0005** | **0.0003** | **0.0003** | **0.0002** | **<0.0001** | 0.14 | 0.81 |
| VacA<sup>-</sup> | 0.78 | 0.78 | 0.78 | 0.78 | 0.59 | 0.78 | 0.78 | 0.78 | 0.59 | **0.0044** |

**Table S4.** Mammalian cell lines and bacterial strains

| Cell lines | Genotype | Phenotype | Source &/or Reference |
| --- | --- | --- | --- |
| AGS | Wild type | n/a | ATCC CRL-1739 |
| HeLa | Wild type | n/a | ATCC CCL-2 |
| HeLa Hexa KO | Inactivating in/del in six Atg8 orthologues (GABARAP, GABARAPL1, MAP1LC3A, MAP1LC3B, MAP1LC3C and GABARAPL2) | Autophagy defect | M. Lazarou; (27) |
| HeLa Vps39 KO | Inactivating insertion in 1<sup>st</sup> exon of *VPS39* | Defective rab5-to-rab7 endosomal transition, and late endosome-lysosome fusion | M. Lazarou; (27) |
| *H. pylori* strains | Genotype | Phenotype | Source&/or Reference |
| 60190wt (*VacA*<sup>+</sup>*Hp*) | Wild type | Vacuolation positive | R. Peek; (42, 63) |
| P12 | Wild type | Vacuolation positive | (64) |
| 60190 *VacA*-null (*VacA*<sup>-</sup>*Hp*) | Insertion of *aphA3* at *vacA* nucleotide 1071 | Inactive truncated *VacA*; vacuolation negative; kanamycin resistant | R. Peek; (65) |
| 60190wt (*VacA*<sup>+</sup>(2)) | Wild type | Vacuolation positive | T. Cover; |
| 60190 Str<sub>312</sub> | Insertion of Strep tag at amino acid 312 | Strep tag II in *VacA* hydrophilic linker; vacuolation positive | T. Cover; (35) |
| 60190Δ6-27 | Deletion of nucleotides encoding *VacA* amino acids 6-27 | Non-channel-forming *VacA*; vacuolation negative | T. Cover; (19) |
| 60190 Str<sub>234</sub> | Insertion of *cat* at 199 bp before *vacA* start codon, Strep tag inserted at amino acid 234 | Strep tag II in *VacA* p31/p28; vacuolation positive; chloramphenicol resistant | This study |
| 60190 Str<sub>525</sub> | Insertion of *cat* at 199 bp before *vacA* start codon, Strep tag inserted at amino acid 525 | Strep tag II in *VacA* p37; vacuolation positive; chloramphenicol resistant | This study |

**Table S5.**
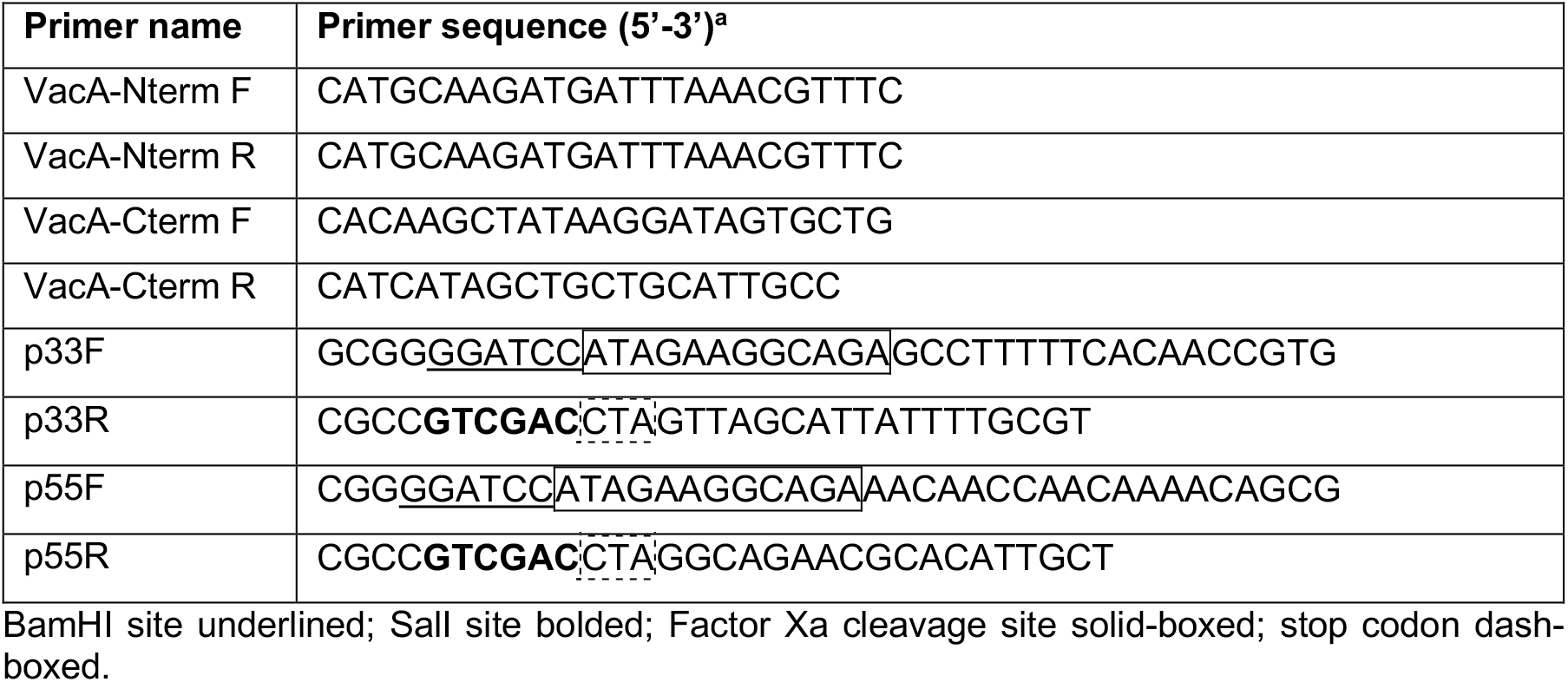
Primer sequences

## References

1. M. M. Coates et al., Burden of non-communicable diseases from infectious causes in 2017: a modelling study. The Lancet. Global health 10.1016/s2214-109x(20)30358-2 (2020).

2. J. G. Kusters, A. H. M. van Vliet, E. J. Kuipers, Pathogenesis of Helicobacter pylori infection. Clin. Microbiol. Rev. 19, 449–490 (2006).

3. B. N. Dong, D. Y. Graham, Helicobacter pylori infection and antibiotic resistance: a WHO high priority? Nature reviews. Gastroenterology & hepatology 14, 383–384 (2017).

4. J. A. Winter et al., A role for the vacuolating cytotoxin, VacA, in colonization and Helicobacter pylori-induced metaplasia in the stomach. J. Infect. Dis. 210, 954–963 (2014).

5. A. Altobelli et al., Helicobacter pylori VacA Targets Myeloid Cells in the Gastric Lamina Propria To Promote Peripherally Induced Regulatory T-Cell Differentiation and Persistent Infection. mBio 10, e00261–00219 (2019).

6. N. R. Salama, G. Otto, L. Tompkins, S. Falkow, Vacuolating cytotoxin of Helicobacter pylori plays a role during colonization in a mouse model of infection. Infect. Immun. 69, 730–736 (2001).

7. M. Oertli et al., Helicobacter pylori γ-glutamyl transpeptidase and vacuolating cytotoxin promote gastric persistence and immune tolerance. Proc. Natl. Acad. Sci. U. S. A. 110, 3047–3052 (2013).

8. K. Zhang et al., Cryo-EM structures of Helicobacter pylori vacuolating cytotoxin A oligomeric assemblies at near-atomic resolution. Proc. Natl. Acad. Sci. U. S. A. 10.1073/pnas.1821959116 (2019).

9. M. Su et al., Cryo-EM Analysis Reveals Structural Basis of Helicobacter pylori VacA Toxin Oligomerization. J. Mol. Biol. 10.1016/j.jmb.2019.03.029 (2019).

10. N. C. Gauthier et al., Helicobacter pylori VacA cytotoxin: a probe for a clathrin-independent and Cdc42-dependent pinocytic pathway routed to late endosomes. Mol. Biol. Cell 16, 4852–4866 (2005).

11. M. de Bernard et al., Helicobacter pylori toxin VacA induces vacuole formation by acting in the cell cytosol. Mol. Microbiol. 26, 665–674 (1997).

12. D. Ye, D. C. Willhite, S. R. Blanke, Identification of the minimal intracellular vacuolating domain of the Helicobacter pylori vacuolating toxin. J. Biol. Chem. 274, 9277–9282 (1999).

13. S. L. Palframan, T. Kwok, K. Gabriel, Vacuolating cytotoxin A (VacA), a key toxin for Helicobacter pylori pathogenesis. Front Cell Infect Microbiol 2 (2012).

14. F. Tombola et al., Helicobacter pylori vacuolating toxin forms anion-selective channels in planar lipid bilayers: possible implications for the mechanism of cellular vacuolation. Biophys. J. 76, 1401–1409 (1999).

15. M. Kimura et al., Vacuolating cytotoxin purified from Helicobacter pylori causes mitochondrial damage in human gastric cells. Microb. Pathog. 26, 45–52 (1999).

16. M. S. McClain, A. C. Beckett, T. L. Cover, Helicobacter pylori Vacuolating Toxin and Gastric Cancer. Toxins (Basel) 9 (2017).

17. I. Szabo et al., Formation of anion-selective channels in the cell plasma membrane by the toxin VacA of Helicobacter pylori is required for its biological activity. EMBO J. 18, 5517–5527 (1999).

18. J. N. Foegeding, R. R. Caston, S. M. McClain, D. M. Ohi, L. T. Cover, An Overview of Helicobacter pylori VacA Toxin Biology. Toxins (Basel) 8 (2016).

19. A. D. Vinion-Dubiel et al., A dominant negative mutant of Helicobacter pylori vacuolating toxin (VacA) inhibits VacA-induced cell vacuolation. J. Biol. Chem. 274, 37736–37742 (1999).

20. C. Genisset et al., A Helicobacter pylori vacuolating toxin mutant that fails to oligomerize has a dominant negative phenotype. Infect. Immun. 74, 1786–1794 (2006).

21. D. M. Czajkowsky, H. Iwamoto, T. L. Cover, Z. Shao, The vacuolating toxin from Helicobacter pylori forms hexameric pores in lipid bilayers at low pH. Proc. Natl. Acad. Sci. U. S. A. 96, 2001–2006 (1999).

22. T. L. Cover, P. I. Hanson, J. E. Heuser, Acid-induced dissociation of VacA, the Helicobacter pylori vacuolating cytotoxin, reveals its pattern of assembly. J. Cell Biol. 138, 759–769 (1997).

23. J. L. Telford et al., Gene structure of the Helicobacter pylori cytotoxin and evidence of its key role in gastric disease. J. Exp. Med. 179, 1653–1658 (1994).

24. D. Burroni et al., Deletion of the major proteolytic site of the Helicobacter pylori cytotoxin does not influence toxin activity but favors assembly of the toxin into hexameric structures. Infect. Immun. 66, 5547–5550 (1998).

25. I.-J. Kim, S. R. Blanke, Remodeling the host environment: modulation of the gastric epithelium by the Helicobacter pylori vacuolating toxin (VacA). Front Cell Infect Microbiol 2, 37–37 (2012).

26. E. J. Gillespie et al., Selective inhibitor of endosomal trafficking pathways exploited by multiple toxins and viruses. Proc. Natl. Acad. Sci. U. S. A. 110, E4904–4912 (2013).

27. T. N. Nguyen et al., Atg8 family LC3/GABARAP proteins are crucial for autophagosome-lysosome fusion but not autophagosome formation during PINK1/Parkin mitophagy and starvation. J. Cell Biol. 215, 857–874 (2016).

28. V. M. Gordon, S. H. Leppla, Proteolytic activation of bacterial toxins: role of bacterial and host cell proteases. Infect. Immun. 62, 333–340 (1994).

29. N. C. Gauthier et al., Early endosomes associated with dynamic F-actin structures are required for late trafficking of H. pylori VacA toxin. J. Cell Biol. 177, 343–354 (2007).

30. S. E. Ivie et al., Helicobacter pylori VacA subdomain required for intracellular toxin activity and assembly of functional oligomeric complexes. Infect. Immun. 76, 2843–2851 (2008).

31. F. Calore et al., Endosome-mitochondria juxtaposition during apoptosis induced by H. pylori VacA. Cell Death Differ. 17, 1707–1716 (2010).

32. H. Iwamoto, D. M. Czajkowsky, T. L. Cover, G. Szabo, Z. Shao, VacA from Helicobacter pylori: a hexameric chloride channel. FEBS Lett. 450, 101–104 (1999).

33. E. H. Heuberger, L. M. Veenhoff, R. H. Duurkens, R. H. Friesen, B. Poolman, Oligomeric state of membrane transport proteins analyzed with blue native electrophoresis and analytical ultracentrifugation. J. Mol. Biol. 317, 591–600 (2002).

34. M. S. McClain et al., Essential role of a GXXXG motif for membrane channel formation by Helicobacter pylori vacuolating toxin. J. Biol. Chem. 278, 12101–12108 (2003).

35. C. Gonzalez-Rivera et al., A Nonoligomerizing Mutant Form of Helicobacter pylori VacA Allows Structural Analysis of the p33 Domain. Infect. Immun. 84, 2662–2670 (2016).

36. B. Geny, M. R. Popoff, Bacterial protein toxins and lipids: pore formation or toxin entry into cells. Biol. Cell. 98, 667–678 (2006).

37. H. Barth, K. Aktories, M. R. Popoff, B. G. Stiles, Binary bacterial toxins: biochemistry, biology, and applications of common Clostridium and Bacillus proteins. Microbiol. Mol. Biol. Rev. 68, 373-402, table of contents (2004).

38. G. Izaguirre, The Proteolytic Regulation of Virus Cell Entry by Furin and Other Proprotein Convertases. Viruses 11 (2019).

39. N. J. Foegeding et al., Intracellular Degradation of Helicobacter pylori VacA Toxin as a Determinant of Gastric Epithelial Cell Viability. Infect. Immun. 87 (2019).

40. T. M. Pyburn et al., Structural organization of membrane-inserted hexamers formed by Helicobacter pylori VacA toxin. Mol. Microbiol. 102, 22–36 (2016).

41. R. D. Leunk, P. T. Johnson, B. C. David, W. G. Kraft, D. R. Morgan, Cytotoxic activity in broth-culture filtrates of Campylobacter pylori. J. Med. Microbiol. 26, 93–99 (1988).

42. T. L. Cover, M. J. Blaser, Purification and characterization of the vacuolating toxin from Helicobacter pylori. J. Biol. Chem. 267, 10570–10575 (1992).

43. T. L. Cover, C. P. Dooley, M. J. Blaser, Characterization of and human serologic response to proteins in Helicobacter pylori broth culture supernatants with vacuolizing cytotoxin activity. Infect. Immun. 58, 603–610 (1990).

44. E. Papini et al., Cellular vacuoles induced by Helicobacter pylori originate from late endosomal compartments. Proc. Natl. Acad. Sci. U. S. A. 91, 9720–9724 (1994).

45. P. Lupetti et al., Oligomeric and subunit structure of the Helicobacter pylori vacuolating cytotoxin. J. Cell Biol. 133, 801–807 (1996).

46. P. Boquet, V. Ricci, Intoxication strategy of Helicobacter pylori VacA toxin. Trends Microbiol. 20, 165–174 (2012).

47. K. A. Gangwer et al., Crystal structure of the Helicobacter pylori vacuolating toxin p55 domain. Proc. Natl. Acad. Sci. U. S. A. 104, 16293–16298 (2007).

48. V. J. Torres, S. E. Ivie, M. S. McClain, T. L. Cover, Functional properties of the p33 and p55 domains of the Helicobacter pylori vacuolating cytotoxin. J. Biol. Chem. 280, 21107–21114 (2005).

49. M. de Bernard et al., Identification of the Helicobacter pylori VacA toxin domain active in the cell cytosol. Infect. Immun. 66, 6014–6016 (1998).

50. G. Domanska et al., Helicobacter pylori VacA toxin/subunit p34: targeting of an anion channel to the inner mitochondrial membrane. PLoS Pathog. 6, e1000878 (2010).

51. O. Odumosu, D. Nicholas, H. Yano, W. Langridge, AB toxins: a paradigm switch from deadly to desirable. Toxins (Basel) 2, 1612–1645 (2010).

52. R. J. Gorrell et al., A novel NOD1- and CagA-independent pathway of interleukin-8 induction mediated by the Helicobacter pylori type IV secretion system. Cell. Microbiol. 15, 554–570 (2013).

53. T. Sato et al., Single Lgr5 stem cells build crypt-villus structures in vitro without a mesenchymal niche. Nature 459, 262–265 (2009).

54. G. A. Reid, G. Schatz, Import of proteins into mitochondria. Yeast cells grown in the presence of carbonyl cyanide m-chlorophenylhydrazone accumulate massive amounts of some mitochondrial precursor polypeptides. J. Biol. Chem. 257, 13056–13061 (1982).

55. I. Gentle, K. Gabriel, P. Beech, R. Waller, T. Lithgow, The Omp85 family of proteins is essential for outer membrane biogenesis in mitochondria and bacteria. J. Cell Biol. 164, 19–24 (2004).

56. H. Alonso et al., Structural and mechanistic insight into alkane hydroxylation by Pseudomonas putida AlkB. Biochem. J. 460, 283–293 (2014).

57. M. Bern, Y. J. Kil, C. Becker, Byonic: advanced peptide and protein identification software. Current Protocols in Bioinformatic Chapter 13, Unit13.20 (2012).

58. E. Gasteiger et al., ExPASy: The proteomics server for in-depth protein knowledge and analysis. Nucleic Acids Res. 31, 3784–3788 (2003).

59. D. C. Montefiori, W. E. Robinson, Jr., S. S. Schuffman, W. M. Mitchell, Evaluation of antiviral drugs and neutralizing antibodies to human immunodeficiency virus by a rapid and sensitive microtiter infection assay. J. Clin. Microbiol. 26, 231–235 (1988).

60. T. L. Cover, W. Puryear, G. I. Perez-Perez, M. J. Blaser, Effect of urease on HeLa cell vacuolation induced by Helicobacter pylori cytotoxin. Infect. Immun. 59, 1264–1270 (1991).

61. E. F. Pettersen et al., UCSF Chimera--a visualization system for exploratory research and analysis. J. Comput. Chem. 25, 1605–1612 (2004).

62. L. A. Kane, M. J. Youngman, R. E. Jensen, J. E. Van Eyk, Phosphorylation of the F(1)F(o) ATP synthase beta subunit: functional and structural consequences assessed in a model system. Circ. Res. 106, 504–513 (2010).

63. M. H. Forsyth, J. C. Atherton, M. J. Blaser, T. L. Cover, Heterogeneity in levels of vacuolating cytotoxin gene (vacA) transcription among Helicobacter pylori strains. Infect. Immun. 66, 3088–3094 (1998).

64. R. Haas, T. F. Meyer, J. P. van Putten, Aflagellated mutants of Helicobacter pylori generated by genetic transformation of naturally competent strains using transposon shuttle mutagenesis. Mol. Microbiol. 8, 753–760 (1993).

65. T. L. Cover, M. K. Tummuru, P. Cao, S. A. Thompson, M. J. Blaser, Divergence of genetic sequences for the vacuolating cytotoxin among Helicobacter pylori strains. J. Biol. Chem. 269, 10566–10573 (1994).

